# Tension at intercellular junctions is necessary for accurate orientation of cell division in the epithelium plane

**DOI:** 10.1101/2022.01.30.478396

**Authors:** Ana Lisica, Jonathan Fouchard, Manasi Kelkar, Tom P. J. Wyatt, Julia Duque, Anne-Betty Ndiaye, Alessandra Bonfanti, Buzz Baum, Alexandre J. Kabla, Guillaume T. Charras

**Affiliations:** London Centre for Nanotechnology, University College London, London, UK; Department of Civil and Environmental Engineering, Politecnico di Milano, Milan, 20133, Italy; MRC-Laboratory for Molecular Cell Biology, University College London, London and MRC-Laboratory of Molecular Biology, Cambridge, UK; Department of Engineering, University of Cambridge, Cambridge, UK; Institute for the Physics of Living Systems, University College London, London, UK; Department of Cell and Developmental Biology, University College London, London, UK

**Keywords:** spindle orientation, tissue tension, out-of-plane division

## Abstract

The direction in which a cell divides is set by the orientation of its mitotic spindle and is important for determining cell fate, controlling tissue shape and maintaining tissue architecture. Division perpendicular to the plane of the substrate can promote tissue stratification during development or wound healing, but also metastasis when orientation is aberrant. Much is known about the molecular mechanisms involved in setting the spindle orientation. However, less is known about the contribution of mechanical factors, such as tissue tension, in ensuring spindle orientation in the plane of the epithelium, despite epithelia being continuously subjected to mechanical stresses. Here, we used suspended epithelial monolayers devoid of extracellular matrix and subjected to varying levels of tissue tension to study the orientation of division relative to the tissue plane. We found that decreasing tissue tension by compressing the monolayers or by inhibiting myosin contractility leads to a higher frequency of out-of-plane divisions. Reciprocally, accurate in-plane division can be restored by increasing tissue tension by increasing cell contractility or by tissue stretching. By considering the full three-dimensional geometry of the epithelium, we show that spindles are sensitive to tissue tension, independently of cell shape, through its impact on the tension at subcellular surfaces. Overall, our data suggest that accurate spindle orientation in the plane of the epithelium necessitates the presence of a sufficiently large tension at intercellular junctions.

**Significance statement:** In growing epithelia, divisions are typically oriented in the plane of the tissue to drive expansion. In some organs, divisions are then re-oriented so that they occur perpendicular to the epithelium plane to drive tissue stratification and cell differentiation. When uncontrolled, this switch in orientation can lead to defects in tissue organisation and, in cancer, likely contribute to metastasis. While much is known about the molecular mechanisms controlling mitotic spindle orientation, less is known about the role of mechanical factors. Here we use mechanical and chemical perturbations to show that mechanics plays a role in controlling the plane of division. Overall, our data suggest that the orientation of spindles in the epithelium plane requires sufficient tension across intercellular junctions.

## Introduction

Orientation of cell division plays a key role in the regulation of tissue growth, cell fate and differentiation during development as well as in adult tissue homeostasis [1, 2, 3]. In monolayered epithelia, divisions typically occur in the plane of the epithelium (XY plane) – driving tissue expansion. In some epithelia, divisions can then be reoriented such that they occur perpendicular to the epithelium plane to drive tissue stratification and cell differentiation. This has been studied in detail in epidermal tissues, where the division of stem cells perpendicular to the plane of the basal layer gives rise to one basal daughter, that retains its stem cell identity, and one suprabasal daughter, that goes on to differentiate and contributes to stratification of the tissue [4, 5]. However, in other contexts, aberrant out-of-plane divisions can lead to failures in morphogenesis [6] and may contribute to cancer metastasis (reviewed in [7, 8]).

The molecular mechanisms and mechanical cues controlling the orientation of cell division within the plane of epithelia have been the focus of much attention. The axis of cell division is set by the orientation of the mitotic spindle, which is controlled in turn by a conserved protein complex composed of Gαi, LGN and NuMA. During mitosis, this complex is localised at the cell cortex where it recruits dynein motors, which exert pulling forces on the astral microtubules resulting in a torque on the spindle. Aside from pulling forces, pushing forces can arise from microtubules polymerising against their site of interaction with the cell cortex and these participate in spindle centring both *in vitro* and *in vivo* [9, 10, 11]. During normal physiological function and development, tissues are continuously subjected to mechanical stress and, consequently, mechanical stresses also participate in regulating the orientation of in-plane cell division [12, 13, 14, 15, 16, 17, 18]. To add further complexity, molecular, geometrical, and mechanical cues appear to interplay to control orientation. Computational studies indicate that the interplay between cellular aspect ratio and cortical pulling forces can control the orientation of the spindle [19]. Recently, the presence of isotropic tissue tension generated by myosin was shown to be necessary to enable spindles to orient toward the long cell axis in cells within the Drosophila notum [16]. Consequently, factors that affect astral microtubules dynamics and stability, classic polarity pathways, cell shape and external forces, all contribute to the regulation of spindle orientation and division (reviewed in [20, 21]).

What factors control the orientation of division out-of-plane is comparatively less well understood and, in particular, little is known about the impact of mechanical forces. In some tissues, the Gαi, LGN and NuMA complex is localised exclusively to intercellular junctions and excluded from the apical domain by aPKC phosphorylation, thus constraining division to the plane of the epithelium [13, 22, 23]. Conversely, in later stages of mouse epidermis morphogenesis or in Drosophila neuroblasts, the relocalisation of LGN to the apical cell surface orients spindles along the apico-basal axis [4, 24, 25, 26]. Interestingly, however, out-of-plane cell division can take place in the absence of the Gαi, LGN and NuMA complex, for example during early development of the mouse epidermis when these proteins are not yet expressed [24]. This has led to the exploration of geometrical cues (such as cell shape and local cell density) as additional factors modulating the early switch from planar to perpendicular divisions. The mouse epidermis starts as a single layer of cells which divide within the plane of the tissue, until an increase in cell density promotes tissue stratification that is driven by a switch in division orientation [27], that has been hypothesised to arise because of a decrease in tissue stress concomitant with the increase in cell density. One challenge in understanding the contribution of mechanical stress to the orientation of division is that direct measurements of cell- and tissue-scale stresses are very difficult. Furthermore, teasing out the relative importance of stress, deformation, and molecular cues is complex because mechanical cues affect cell shape and protein localization. Indeed, decreases in tissue tension have been reported to reduce LGN and E-Cadherin signals at cell-cell contacts [27, 14]. In summary, it remains unclear what exact stimulus or combination of stimuli influence the plane of cell division.

To investigate the relative contribution of geometrical and mechanical cues in regulating out-of-plane spindle orientation, we imaged cell divisions in suspended epithelial monolayers. We show that, when cells are exposed to a moderate level of tissue tension, divisions are robustly oriented within the plane of the monolayer. Strikingly, however, a decrease in tissue tension induced by chemical treatment or by compressing the tissue increased the frequency of out-of-plane divisions. Our data indicate that tension at intercellular junctions is required to enable cortical regulators to efficiently orient division in the plane of the epithelium – revealing a role for the subcellular distribution of cell stress in this process.

## Results

### Application of uniaxial compressive strain promotes division out-of-plane

In our experiments, we used epithelial MDCK monolayers devoid of a substrate and suspended between test rods (**Fig 1A, B**, **Methods**, [28, 29]). This experimental system allows the accurate control of tissue-scale stress and strain, while simultaneously allowing for imaging of the subcellular localization of proteins and cell shape.

**Figure 1.**
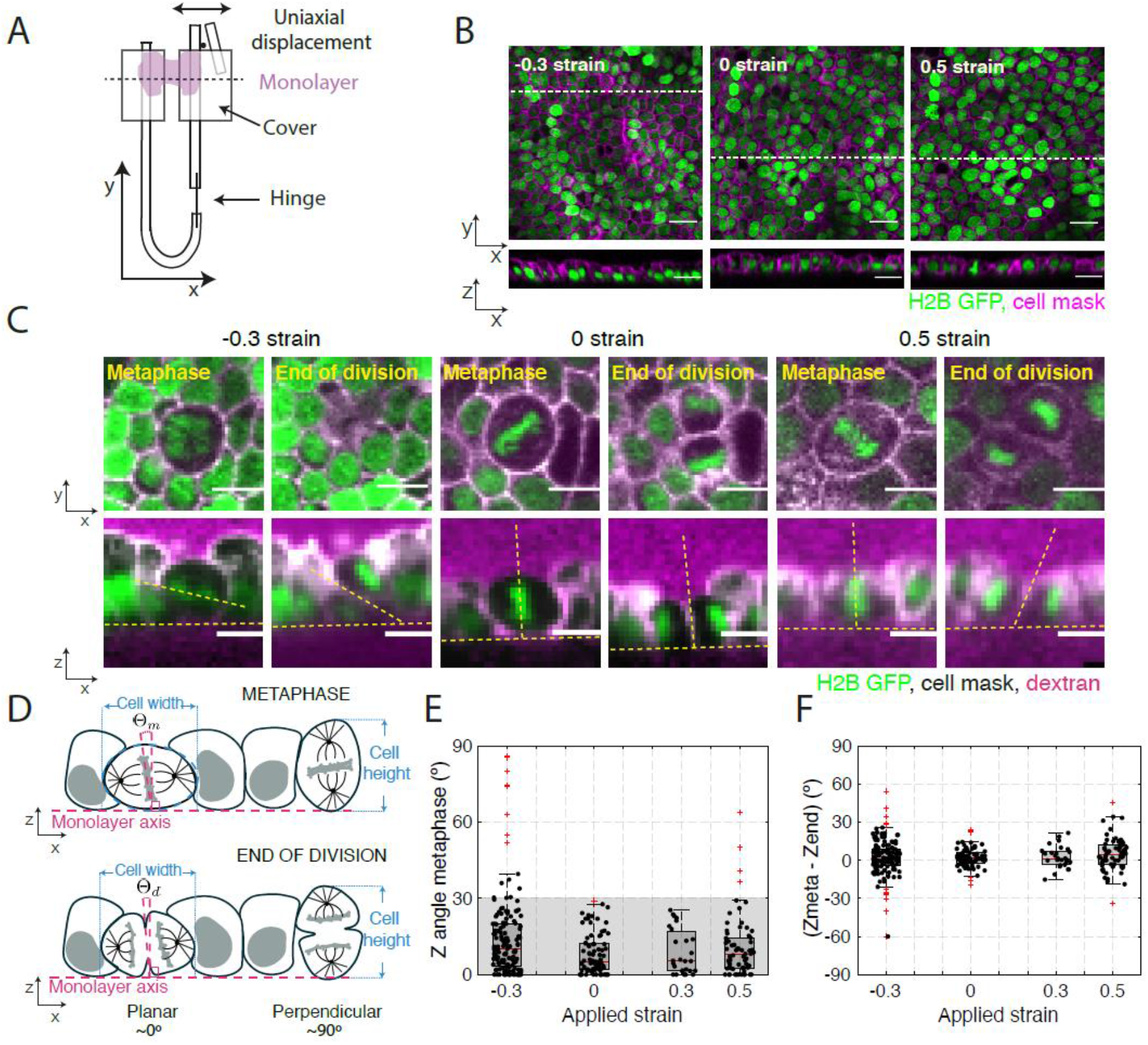
Application of uniaxial compressive strain to epithelial monolayers promotes division out-of-plane. **A.** Diagram of the device for mechanical manipulation of suspended MDCK monolayers. The U-shaped device consists of a rigid arm and a flexible arm. Small coverglasses (grey) are glued at the extremities of each arm creating a gap of ^~^500μm. A drop of collagen is then polymerised in this gap and cells are seeded on top of it. Once the cells form a monolayer spanning the gap (magenta), the collagen is removed by enzymatic digestion leaving the monolayer suspended between the top plates. Uniaxial strain can be applied to the monolayer by displacing the flexible arm with a motorised manipulator. **B.** Representative images of suspended MDCK cell monolayers subjected different strain conditions in the XY (top) and XZ (bottom) planes. The strain to which the monolayer was subjected is indicated in the top left corner. Nuclei are marked with H2B GFP (green) and cell membrane with CellMask (magenta). Dashed white lines indicate the planes at which the XZ profiles were taken. Scale bar: 10 μm. **C.** Examples of cell divisions in MDCK monolayers subjected to different strains viewed in the XY (top) and XZ (bottom) plane. Each cell is shown at metaphase and at the end of cytokinesis. Nuclei are marked with H2B GFP (green), cell membranes are visualised with CellMask (white) and Alexa Fluor 647 dextran is added to the medium to allow visualization of the cell outlines. In the profile views, the horizontal yellow dashed lines indicate the plane of the monolayer, while the slanted and vertical dashed lines indicate the orientation of the metaphase plate or the division furrow. Scale bar: 10 μm. **D.** Diagram of the spindle and division orientation measurements in the XZ plane. The spindle Z angle at metaphase (Θ_*m*_) was calculated as the angle between the line passing through the metaphase plate and the line perpendicular to the monolayer axis. Similarly, the spindle Z angle at the end of division (Θ_*d*_) was calculated as the angle between the line passing through the closed cytokinetic furrow and the line perpendicular to the monolayer axis. **E.** Distribution of spindle Z angles at metaphase (Θ_*m*_) for different applied strains. Gray box highlights Z angles < 30º. The number of mitotic cells examined for each condition was N=147 for −30% strain, N=81 for 0% strain, N=27 for 30% strain, and N=68 for 50% strain. Experiments were performed on n=14 independent days for −30% strain, n=8 independent days for 0% strain, n=4 independent days for 30% strain, and n=8 independent days for 50% strain. **F.** Difference between spindle Z-angles at the beginning of metaphase and the end of division for each applied strain. The data correspond to the same experiments as in E. (**E-F**) Box plots indicate the 25th and 75th percentiles, the red line indicates the median, and the whiskers extend to the most extreme data points that are not outliers. Individual data points are indicated by black dots and outliers by red crosses.

To characterise the orientation of division in suspended epithelia, we acquired confocal stacks of the monolayers every minute for 1 hour and quantified the orientation of cell divisions. To allow visualisation, we imaged cells expressing the nuclear marker H2B-GFP with their cell membranes fluorescently labelled. Orientation of cell divisions was characterised by measuring mitotic spindle orientation relative to the plane of the tissue (or “Z-angle”). Depending on the stage of mitosis, the z-angle was defined as either the angle between the line going through the metaphase plate or the line going through the closed cytokinetic furrow and the line perpendicular to the monolayer plane (see **Methods**, **Fig 1C, D**). We found that in non-perturbed monolayers (exposed to 0% strain), divisions are oriented so that all cells divide within 30° of the monolayer plane (median=5.1°, N=81 cells, **Fig 1E**, **Fig S1A**). Although some cells experienced large transient changes in the spindle Z-angle between metaphase and the end of division, the difference in angle between these two time-points was close to zero (median=1.5°, **Fig 1F**). Therefore, under control conditions, cell divisions occur within the plane of MDCK suspended epithelia along a direction that is set by the orientation of the metaphase spindle, consistent with previous reports examining tissues growing on a substrate [30,31].

Out-of-plane division in epithelia has been observed when cell density is high in physiological and pathological conditions. To mimic this, we subjected epithelial monolayers to a −30% compressive strain that greatly increases cell density while preserving tissue planarity [32] (see **Methods**, **Fig 1B, C**). When we examined the impact on division orientation, we found that compressive strain significantly changed the spindle Z-angle distribution resulting in a doubling of its median (median=10.1°, p=0.003, Wilcoxon rank sum test (WRST)). In contrast, application of tensile strains of 30 and 50% did not significantly change the Z-angle distribution (median=5.3°, p=0.93 and median=8.2°, p=0.24, respectively, WRST, **Fig 1E, S1A**). To compare the prevalence of out-of-plane divisions in each condition, we categorised our data into spindles dividing in the plane (Z-angle<30°) and out-of-plane (Z-angle>30°), and compared conditions using Fischer’s exact test (FET). Whereas no mitotic cell had a Z-angle larger than 30° at metaphase at 0% strain (0/81 divisions), 11% were misoriented when the monolayer was subjected to −30% compressive strain (16/147 divisions). Only compressive strain significantly increased the prevalence of out-of-plane division (p<0.01 for −30% compressive strain; 0/27 divisions, p=1 for 30% strain; 4/68 divisions, p=0.04 for 50% strain; FET). Thus, our results show that application of compressive strain, but not tensile strain, leads to an increase in the frequency of division out-of-plane.

### Changes in cell shape, height, and density do not correlate with more frequent out-of-plane division

As previous work has highlighted roles for cell shape, cell height and cell density in regulating the orientation of division out-of-plane, we examined the relevance of these parameters to the increased out-of-plane division observed in compressed monolayers.

For division in the plane of the tissue to be possible, the dimension of the cell in the XY plane must be larger than the spindle length and the cell height must be larger than the height of chromosomes congregating at the metaphase plate. Compression of epithelial monolayers shortens cells along the axis of compression (X-axis) and lengthens them along the out-of-plane axis (Z-axis) [32]. Conversely, stretch elongates cells along the X-axis and thins them along the Z-axis. Neither manipulation changes the cell size along the Y-axis [32]. Therefore, we compared the cell length along the X-axis to the pole-to-pole length of fluorescently labelled metaphase spindles. The length of the metaphase plate measured from the H2B GFP signal was compared to the cell height in XZ plane (see **Methods**, **Fig 2A**). Our results show that the ratio of cell length to spindle length increased with increasing strain, while the ratio of cell height to metaphase plate length decreased with increasing strain, as expected. However, in all conditions, both ratios remained above 1 (**Fig 2B, C**), indicating that the cell dimensions likely do not constrain spindle movement and orientation.

**Figure 2.**
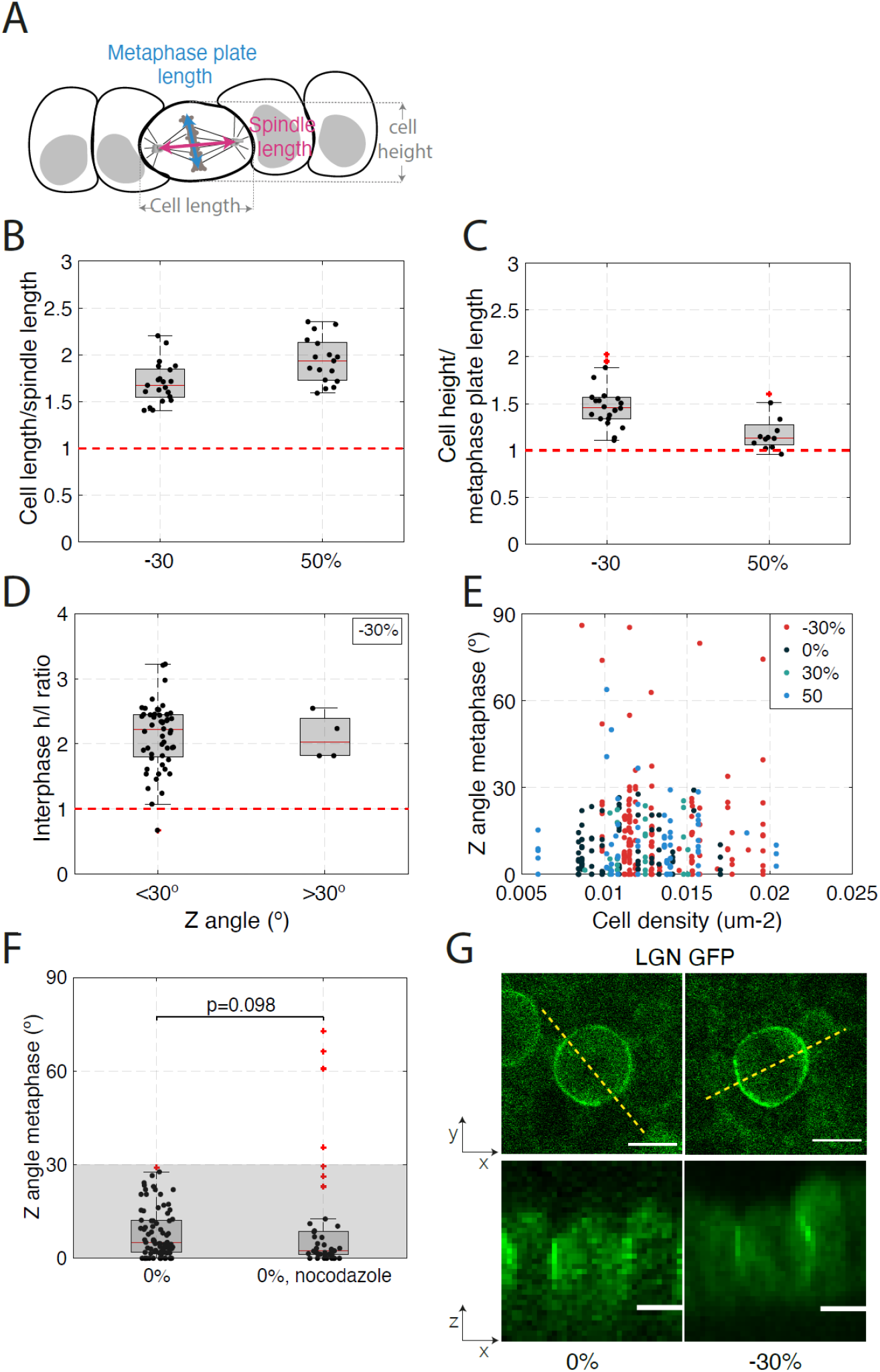
Changes in cell shape, cell density or the spindle positioning machinery do not correlate with increased out-of-plane division induced by mechanical manipulation. (**B-D, F**) Box plots indicate the 25th and 75th percentiles, the red line indicates the median, and the whiskers extend to the most extreme data points that are not outliers. Individual data points are indicated by black dots and outliers by red crosses. **A.** Diagram indicating cell and spindle measurements in the XZ plane. Cell width is measured along the X-axis and cell height along the Z-axis. The bounds of the cell were determined from the CellMask fluorescence signal. The metaphase plate length was determined as the extent of the H2B GFP signal and the spindle length was determined as the pole to pole distance visualised using SiR-tubulin fluorescence signal. **B.** Ratio of cell width to spindle length for dividing cells in compressed (−30%) and stretched monolayers (50%). N=21 cells for −30% strain and N=17 cells for 50% strain. Data from n=2 and n=1 independent days, respectively. **C.** Ratio of cell height to metaphase plate length for dividing cells in compressed (−30%) and stretched monolayers (50%). N=22 cells for −30% strain and N=12 cells for 50% strain. Data from n=2 and n=1 independent days, respectively. **D.** Ratio of cell height *h* to length *l* at interphase as a function of spindle Z angle at metaphase for dividing cells in compressed monolayers (−30%). Metaphase spindle Z-angles were categorised as either in-plane (Θ<30°), or out-of-plane (Θ >30°). N=54 cells from n=14 independent days. **E.** Spindle Z angle as a function of cell density for monolayers subjected to different amplitudes of uniaxial strain. N= 147 cells for −30%, 81 cells for 0%, 27 cells for 30% and 68 cells for 50% strain. Experiments were performed on n=14, n=8, n=4, and n=8 independent days. **F.** Distribution of spindle Z angles at metaphase for untreated monolayers and monolayers treated with 20 nM nocodazole. Wilcoxon rank-sum test (WRST, p=0.098). Gray box highlights Z angles < 30º. N=81 mitotic cells for 0% strain, N=39 for nocodazole treatment. Experiments were performed on n=8 and n=2 independent days, respectively. **G.** Representative localisation of LGN in dividing cells in a monolayer subjected to 0 % and −30 % compressive strain viewed in XY (top) and XZ (bottom) planes. Dashed white lines indicate the locations at which XZ profiles were taken. Scale bars: 10 μm.

Next, we investigated if changes in cell shape might be responsible for the increased incidence of out-of-plane division in compressed monolayers. We characterized cell shape by computing each cell’s height/length (*h/l*) ratio. Spindle Z-angles at metaphase measured in the compressed monolayers (−30%) were categorised as either in-plane (Z-angle <30°) or out-of-plane (Z-angle >30°). We compared *h/l* ratios for cells from each category at interphase and metaphase (**Fig 2D**, **Fig S2A**). Our data show no difference between categories, suggesting that the higher frequency of out-of-plane spindle orientation observed in compressed monolayers is not correlated with changes in cell shape.

Finally, we examined the impact of cell density changes induced by deformation of monolayers. Previous work has demonstrated that epithelia possess a well-defined homeostatic density that they strive to return to following perturbation [33, 34]. In compressed monolayers, out-of-plane divisions may therefore act to prevent further increases in cell density. In suspended MDCK monolayers, density can only be varied between 0.005-0.02 cells.μm^-2^ because less dense monolayers rupture, while denser seeding leads to ‘lumpy’ monolayers with regions of multi-layering. In the experimentally attainable range of cell densities, our results show no correlation between cell density and the metaphase spindle Z-angle (**Fig 2E**). Overall, our data indicate that changes in cell shape, height, or density do not correlate with increased frequency of out-of-plane division.

### Interaction between astral microtubules and cortical regulators is necessary for orientation of division in the plane

Spindle orientation results from interactions of astral microtubules with the actin cortex and previous work has shown that loss of astral microtubules results in an increase in spindle Z-angle in isolated MDCK cells on a substrate [31]. Therefore, we treated suspended monolayers, not subjected to strain, with a concentration of nocodazole sufficiently low to affect astral microtubules without preventing division [12] (**Fig S2B**). Nocodazole treatment resulted in an increase in the incidence of spindle Z-angles larger than 30° at metaphase (5/43 divisions, p=0.004, FET, **Fig 2F**). This suggested that interaction between astral microtubules and the cortex remains essential for correct spindle orientation despite the lack of a substrate.

Astral microtubules interact with the Gαi, LGN and NuMA complex at the cell membrane. Therefore, we examined the localisation of this complex in suspended monolayers using LGN-GFP as a proxy (see **Methods**). When monolayers were not subjected to strain, LGN localized to the cell periphery in the XY plane at metaphase, with a weaker localization at the cell equator in the vicinity of the metaphase plate, and stronger localisation at the poles. In the XZ plane, LGN localized homogenously along the entire height of the intercellular junction, consistent with previous reports examining MDCK cysts [13, 23] (**Fig 2G**). Importantly, the application of a 30% compressive strain did not change the localization of LGN in either the XY or XZ planes. These data show that, while division in the XZ plane requires interactions between astral microtubules and the cortex, the increase in out-of-plane cell divisions in response to compressive strain does not correlate with a change in the localisation of cortical regulators at the cell periphery.

### Reduction of tissue tension increases the frequency of out-of-plane division

Previous work has shown that tissue tension affects spindle orientation in the XY plane of the monolayer [2, 17, 18, 15, 16] and other work hypothesised that it may influence spindle positioning in the XZ plane in the mouse epidermis [24, 27]. However, an accurate characterization of tissue stresses and how they influence spindle orientation is missing because of the difficulty of measuring stress *in vivo*. Further complexity arises because, in living tissues, tension can emerge from either active or passive processes. Active stress originates from the action of myosin motors on the cytoskeleton that is transmitted to other cells through adhesion complexes to generate tissue tension; whereas passive stresses arise from deformation of cytoskeletal networks in response to external forces applied on the tissue. Previous work has shown that both types of stress are present in epithelial monolayers subjected to deformation [35]. Therefore, we examined the influence of active and passive stresses on cell division orientation.

We first characterized tissue tension in experiments in which monolayers were subjected to mechanical manipulations (see **Methods**, **Fig 3A**, [32]). At 0% strain, the tissue tension was 238±148 Pa (**Fig 3C, D**). The application of −30% compressive strain significantly reduced tissue tension to 52±51 Pa (p= 0.004, WRST compared to 0% strain), suggesting that the frequency of out-of-plane division may increase when tissue tension is low.

**Figure 3.**
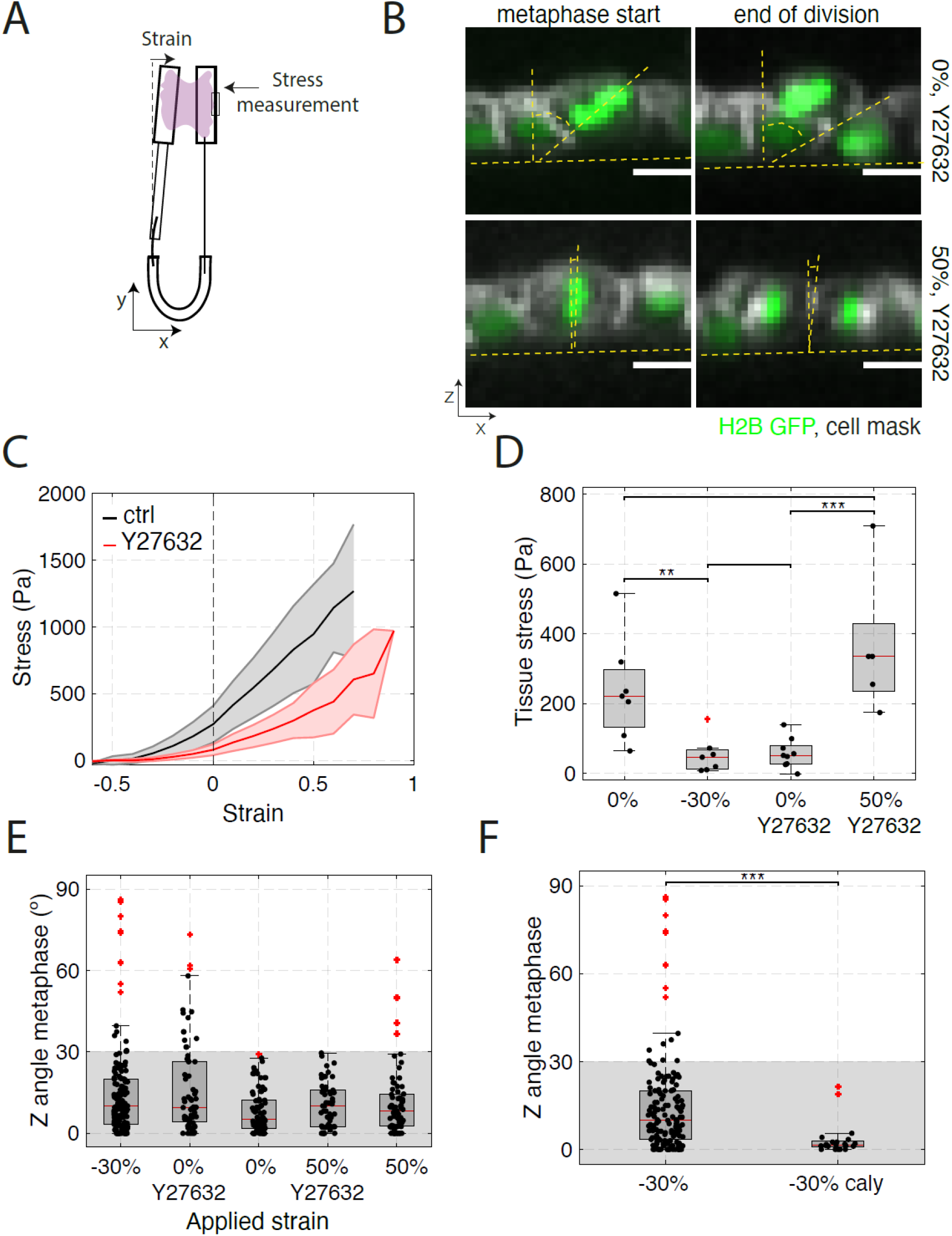
The accuracy of in-plane cell division is controlled by tissue tension. (**C, E, F**) Box plots indicate the 25th and 75th percentiles, the red line indicates the median, and the whiskers extend to the most extreme data points that are not outliers. Individual data points are indicated by black dots and outliers by red crosses. **A.** Diagram of the device used for measurement of stress during application of uniaxial strain. Monolayers are cultured between a reference rod and a flexible rod. Measurement of the deflection of the flexible rod allows determination of the stress applied on the epithelium (**see methods**). **B.** Example of cell division in a monolayer treated with 50 μM Y27632 at 0% (top row) and 50% strain (bottom row). Spindle Z angle is measured at the beginning of metaphase (left) and at the end of division (right). Nucleic acids are visualised with H2B GFP (green), the cell membrane is labelled with CellMask 568 dye (white). In the profile views, the horizontal yellow dashed lines indicate the plane of the monolayer, while the slanted and vertical dashed lines indicate the orientation of the metaphase plate or the division furrow. Scale bars: 10 μm. **C.** Tissue stress as a function of strain in response to a ramp in deformation applied at low strain rate (0.5%.s^-1^) for control (black) and Y27632-treated (red) monolayers. Solid lines indicate the mean and the shaded area shows the standard deviation. N=7 monolayers for control and N=8 monolayers for Y27632 treatment. n= 4 and n=5 independent days, respectively. **D.** Tissue stress in MDCK monolayers as a function of strain in control conditions and in monolayers treated with 50 μM Y27632. WRST, p = 0.0041 (−30%, 0%), p = 0.6255 (0%, 0% + Y27632), p=0.0009 (0% + Y27632, 50% + Y27632), p = 0.1869 (0%, 50% + Y27632). Data from C. **E.** Distribution of spindle Z angle at metaphase for non-treated monolayers at −30 %, 0 % and 50 % strain, and monolayers treated with 50 μM Y27632 at 0 % and 50 % strain. Gray box highlights Z angles < 30º. The number of mitotic cells examined for each condition was N=147 for −30% strain, N=66 for 0% strain with Y27632, N=81 for 0% strain, N=53 for 50% strain with Y27632 and N=68 for 50% strain. Data from n=14, n=8, n=8, n=11, and n=8 independent days, respectively. **F.** Distribution of the spindle Z angles at metaphase for monolayers subjected to −30% strain in control conditions or treated with 35 nM calyculin. WRST, p = 1.945 10^-5^. Gray box highlights Z angles < 30º. N=147 mitotic cells for −30% strain and N=21 for −30% strain with calyculin treatment. Data from n=14 and n=2 independent days, respectively.

Our previous work has shown that, at 0% strain, most of the tension in the tissue originates from active stresses due to myosin contractility in the submembranous actin cortex [32, 35]. Furthermore, in dividing cells, cortical actomyosin plays a crucial role in enabling proper orientation and centring of the mitotic spindle [36, 37, 16]. This led us to investigate whether a reduction of tissue stress induced through the treatment of monolayers with an inhibitor of Rho-kinase would also increase the frequency of out-of-plane divisions in the absence of tissue compression. Treatment with Y27632 significantly decreased tissue tension to 57±42 Pa at 0% strain, similar to the effect of a −30% compression (**Fig 3B, C**, **D**, p=0.12 compared to −30% compression, WRST). As in compressed monolayers, divisions in Y27632-treated monolayers exhibited increased frequency of out-of-plane division (**Fig 3B, E**, 14/66 divisions, p=0.001, FET). Again, we found no correlation with cell shape descriptors (*h/l* ratio) for spindles oriented in- or out-of-plane (**Fig S3A, B**). Thus, we observed increased frequency of out-of-plane division in both compressed and Y27632-treated monolayers despite cell shape being different between these two conditions (**Fig S3C, D**). Again, the localisation of LGN in the XY and XY planes was not affected by Y27632 treatment (**Fig S3G**). Overall, these data further suggest that changes in shape, density, or localisation of pulling forces cannot account for the observed increase in the frequency of out-of-plane division but that a decrease in tissue tension might.

### Increasing active or passive stress reduces out-of-plane divisions in monolayers with low tension

Since decreasing tissue tension through chemical treatment or mechanical manipulations increased out-of-plane divisions, we investigated whether the reduced accuracy of in-plane orientation of division could be rescued by increasing tissue tension by orthogonal means.

To do so, we first examined if the increased frequency of out-of-plane division induced by chemical inhibition of contractility could be rescued by stretching the monolayer such that the original tissue tension was restored. For this, we characterised the stress response of Y27632-treated monolayers and compared their stress-strain curves to those of control monolayers. Our measurements showed that application of a 50% stretch to a Y27632-treated monolayer results in a tissue tension comparable to that in control monolayers at 0% strain (**Fig 3C, D**). We then examined the orientation of cell division in monolayers treated with Y27632 subjected to a 50% stretch. In these conditions, spindle Z-angles returned to a distribution similar to that in non-stretched control monolayers (**Fig 3E**, p=0.12, WRST) and the frequency of out-of-plane division returned to control values (1/53 divisions, p=0.002 FET compared to Y27632 alone). This was the case despite profound differences in interphase and mitotic cell shape between non-stretched control monolayers and stretched, Y27632-treated monolayers (**Fig S3E, F**), again supporting the idea that cell shape does not influence the frequency of out-of-plane divisions.

Next, we determined if more frequent out-of-plane division orientation induced by a decrease in tension due to compression could be rescued by increasing cell contractility by treating monolayers with calyculin, a phosphatase inhibitor that leads to a 1.5 fold increase in tissue tension in compressed monolayers [32]. We performed experiments in which we compressed monolayers to −30% strain before adding calyculin. Although cytokinesis was perturbed in a fraction of cells, the cells that did undergo cytokinesis divided within the plane of the monolayer as they do in control non-stretched monolayers (0/21 divisions). As a result, the distribution of Z-angles was significantly different to that observed in non-treated, compressed monolayers (**Fig 3F**, p=2×10^-5^, WRST). Importantly, this rescue could be achieved without a significant change in cell shape (**Fig S3H, I**).

Together, these experiments show that an increase in tissue tension is sufficient to reduce the frequency of out-of-plane divisions induced by compression or inhibition of contractility. Remarkably, active and passive tissue stresses appear interchangeable in this regard, which suggests that they are both sensed in the same way by dividing cells.

### Mechanical and chemical manipulations of the tissue have similar effects on the mechanical environment of interphase and mitotic cells

So far, our experiments indicate that more frequent out-of-plane division is associated with low tissue tension but, at the cellular-scale, we do not know if treatments have similar effects on interphase and mitotic cells. Indeed, as suspended MDCK monolayers are primarily composed of cells in interphase with only about 1-2% mitotic cells at any given time, the mechanics of monolayers largely reflects the mechanics of interphase cells, which is controlled by the submembranous actin cortex [35]. However, isolated mitotic cells are mechanically distinct from interphase cells with a higher cortical tension [38, 39, 40], as a result mitotic cells are stiffer than interphase cells and deform less when the tissue is subjected to stretch. Since we hypothesise that mitotic cells respond to tissue tension, we sought to determine if pharmacological treatments and changes in the tissue tension differentially affect mitotic cells and their interphase neighbours.

To test this idea, we first confirmed that the previously reported differences in mechanics between mitotic and interphase cells are also observed in suspended epithelia. To do so, we subjected epithelia to cyclic uniaxial deformation with a 50% amplitude (see **Methods**) and imaged the change in length of interphase cells and mitotic cells along the direction of stretch. Our data showed that interphase cells were ^~^2.5 fold more deformed than mitotic cells (**Fig 4A, B**), indicating that mitotic cells are ^~^2.5 fold more stiff than their interphase neighbours – comparable to the increase in cortical tension noted in isolated mitotic cells.

**Figure 4:**
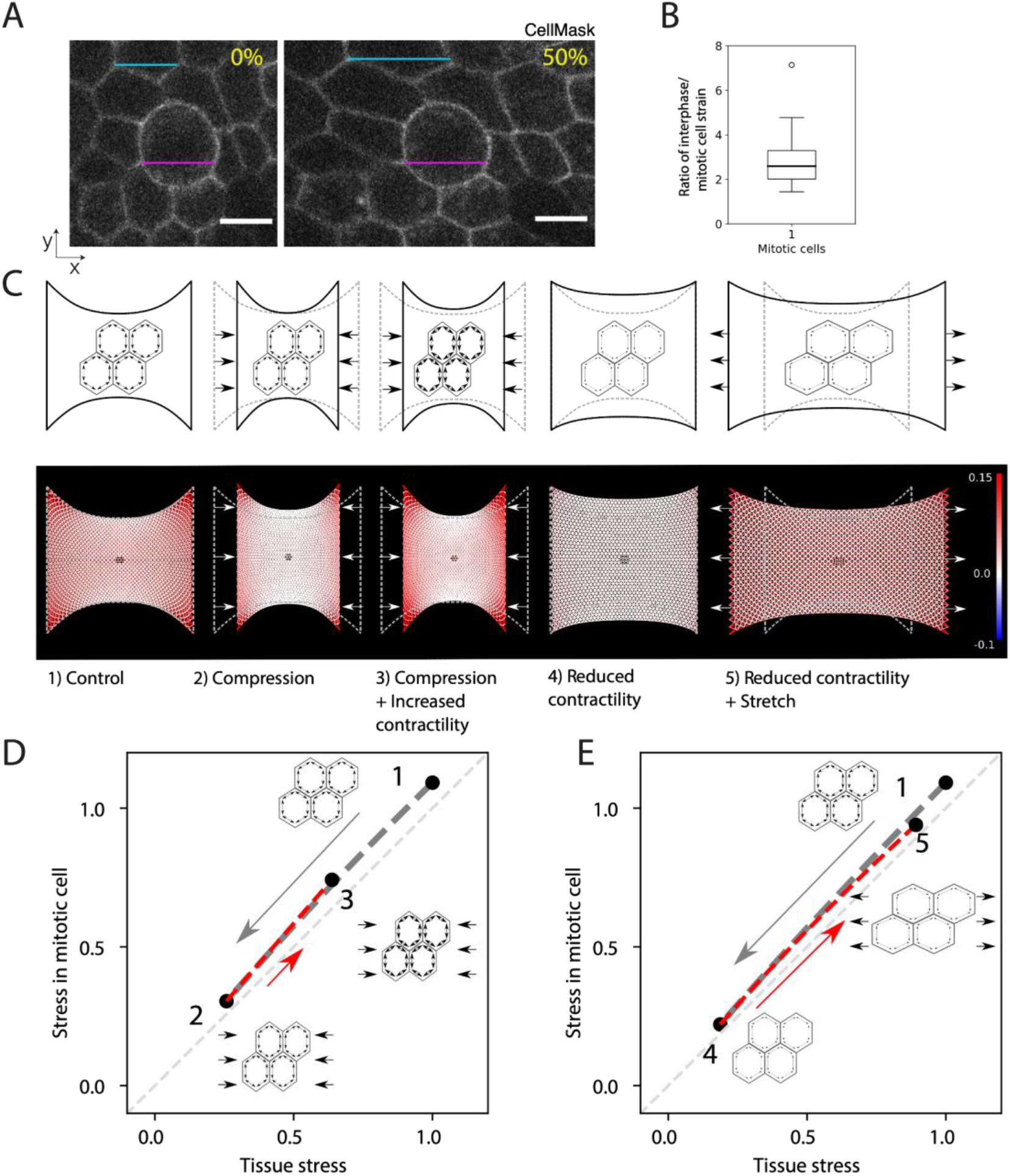
Mechanical stress at the tissue-scale reflects the mechanical environment of interphase and mitotic cells in response to chemical and mechanical manipulations. **A.** Representative images of mitotic cells in a monolayer subjected to 0% strain and 50% strain. The magenta lines denote the length of the mitotic cell in both images, while the blue lines show the length of an interphase cell in both conditions. Cells are labelled with CellMask membrane stain. Scale bars: 10 μm. **B.** Ratio of interphase cell strain to mitotic cell strain, calculated from the cyclical stretching experiments with an amplitude of ^~^50% shown in Fig S4A. Strain is calculated from measurements of the bounding box of the respective cell before and after a stretch is applied to the monolayer. 11 mitotic cells were measured from 11 different monolayers. Mean strain in interphase cells is calculated from the strain in three interphase neighbouring cells that do not have any junction in common with the mitotic cell of interest. The distribution’s median, first and third quartile and range are represented by the central bar, bounding box and whiskers, respectively. **C.** Finite element model predictions of the stress distributions in monolayers subjected to mechanical and chemical manipulations. Top: schematic representation of the experimental conditions: (1) a monolayer in its initial configuration clamped at both ends (0% strain) and subjected to a tensile stress arising due to cell contractility; (2) a monolayer subjected to a −30% compressive strain applied by displacing the test rods; (3) a monolayer subjected to a −30% compressive strain and treated with a drug increasing cell contractility (calyculin); (4) a monolayer at 0% strain treated with a drug decreasing cell contractility (Y27632); (5) a monolayer treated with a drug decreasing cell contractility and subjected to a 50% tensile strain. Bottom: Stress distribution within the monolayer. The mesh representing the monolayer is colour coded to display the stress in each element with red colours representing tensile stress, blue colours compressive stress, and white colours regions of low stress. A mitotic cell simulated as a stiffer inclusion is present in the centre of the monolayer (dark region). Stress distributions are presented for the experimental conditions depicted above. **D.** Stress in a mitotic cell in the centre of a monolayer as a function of the stress in the tissue in response to compressive strain followed by a chemical treatment to increase contractility. In the monolayer’s initial configuration, the stress on the mitotic cell is approximately equal to the tissue stress (state 1, corresponding to condition 1 in panel A). When a compressive strain is applied to the monolayer, the stress in the mitotic cell decreases in proportion to the decrease in the tissue stress (grey dashed line, transition from state 1 to state 2). Increasing prestress in the cells by increasing myosin contractility by calyculin treatment leads to a partial recovery of the tensile stress in the tissue and the mitotic cell (red dashed line, transition from state 2 to state 3). All stresses are normalised to the tissue stress in state 1. **E.** Stress in a mitotic cell in the centre of a monolayer as a function of the stress in the tissue in response to a chemical treatment to decrease contractility followed by application of a tensile strain. In the monolayer’s initial configuration, the stress on the mitotic cell is approximately equal to the tissue stress (state 1, corresponding to condition 1 in panel A). When a chemical treatment that decreases cellular prestress is applied, the stress in the tissue and the cell drops to values close to 0 (grey dashed line, transition to state 1 to state 4). To recover a stress similar to condition 1, a 50% stretch is applied to the monolayer (red dashed line, transition from state 4 to state 5). All stresses are normalised to the tissue stress in state 1.

The larger stiffness of mitotic cells and the smaller deformations that they experience during tissue manipulations suggest that stress may be unevenly distributed around the dividing cell and that they may experience significantly different mechanical stress than the tissue tension. To gain a conceptual understanding of how the tension in mitotic cells evolves relative to the tension in interphase cells and in the tissue, we devised a simple computational model of the monolayer to reproduce the range of experimental conditions studied above and characterize the distribution of stress in the vicinity of a mitotic cell.

In the model, the monolayer is a 2D elastic material discretised into a triangular mesh in which cells are represented as hexagons [41] (**Fig S4**). Because most of the tension is born by the basal layer through the whole tissue, all springs corresponding to interphase cell have the same spring constant *k*. An active tension *γ* is imposed by introducing a difference between the reference length *l_init_* of the junction and its rest length *l*_0_ (see **Materials and Methods**). First, we adjusted the parameters of the model to obtain a stress-strain relationship for the virtual tissue similar to that measured experimentally in suspended monolayers at low strain rates (**Fig 3C**). Then, we modelled the mitotic cell as an inclusion in the centre of the sheet of cells (blue cell, **Fig S4**) with different values for the spring constant *k* and rest length *l*_0_ such that their active tension is 2.5 times larger than in interphase cells [38] and their stiffness increased by a factor 2.5 to fit the lower deformation of mitotic cells observed in response to stretch (**Fig S4A-C**).

Using the model, we can compare the tension in a mitotic cell, with the tension in neighbouring interphase cells and with the overall tissue tension. To simulate mechanical manipulations, we apply a compressive or tensile strain onto the network and to simulate chemical treatments affecting contractility, we change the junctional tension *γ* (**Fig 4C-E**). For all experimental conditions, our simulation data shows that the tension in the mitotic cells remains close to tension in the neighbouring interphase cells and in the tissue as a whole (**Fig 4D, E**). However, the increased rigidity of the mitotic cells slightly amplifies the mechanical stress they experience, despite being less deformed (**Fig 4D, E**). The model also illustrates the mechanical interplay between contractility and extrinsic deformations in controlling the stress to which mitotic cells are subjected. When monolayers are compressed, tissue tension decreases, and as a consequence the stress that the mitotic cells experience also decreases (transition from 1 to 2, **Fig 4D**). With the addition of calyculin to increase contractility, there is an increase in tissue tension as well as in the tension experienced by mitotic cells (transition from 2 to 3, **Fig 4D**). Further, our analysis reveals that the interplay between the increased contractility at the cell periphery and the tissue boundary conditions restores a cell stress tensor very similar to that observed in control conditions (**Fig S5, S6**). Similarly, when contractility was inhibited (transition from 1 to 4, **Fig 4E**), the stress experienced by mitotic cells decreased in proportion to the tissue stress and the stress tensor in mitotic cells became similar to that in compressed monolayers (**Fig S6**). The application of stretch restored the stress experienced by mitotic cells in the model to a level and tensor similar to that in control conditions (transition from 4 to 5, **Fig 4F**, and **Fig S7**). Overall, this analysis leads us to conclude that the tissue tension measured in our experiments is a good proxy for the tension experienced by the mitotic cells.

### Out-of-plane divisions are not linked to mechanical changes specific to mitotic cells

Because cortical mechanics likely play an important role in the mechanical response of cells within the monolayer, we characterized the interaction of mitotic cells with their interphase neighbours by measuring their apical angle of contact Θ_*a,mi*_ (**Fig 5A**). This angle is determined by the balance of tensions in the apical cortices of the mitotic cell and its interphase neighbours, together with the tension at the intercellular junction through the Young-Dupré relationship [42] (see **Methods**). A change in the apical angle of contact Θ_*a,mi*_ in response to a treatment would indicate that the cortical tension in mitotic cells and interphase cells respond differently to treatment. We compared Θ_*a,mi*_ in monolayers treated with DMSO and Y27632, since Rho-kinase inhibition leads to more frequent out-of-plane divisions without any detectable change in mitotic cell shape. Θ_*a,mi*_ did not change in response to treatment (**Fig 5B**), indicating that Rho-kinase inhibition affects cortical tension in a similar way in both mitotic and interphase cells in line with the conclusions of our simulation.

**Figure 5.**
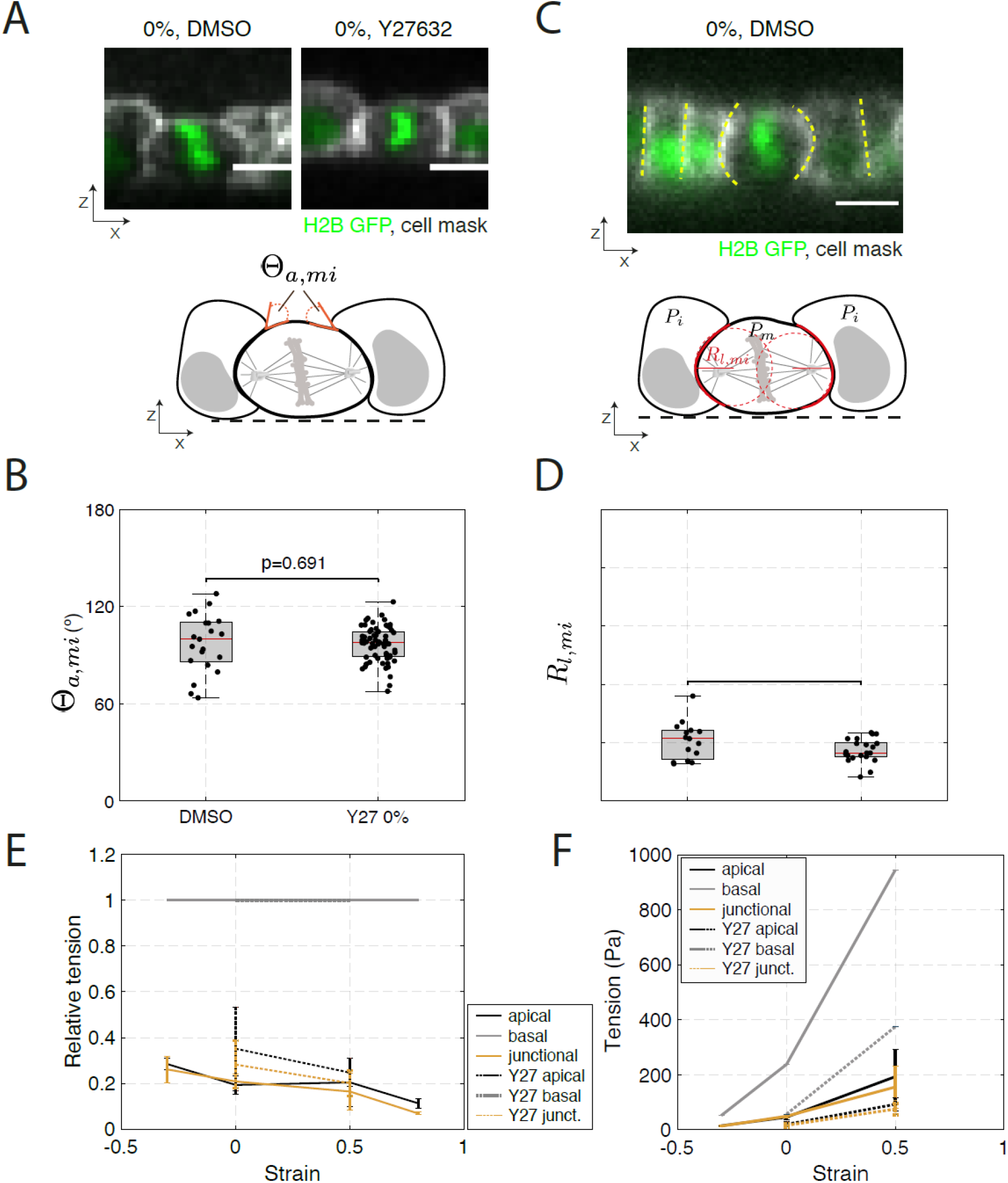
Treatments that decrease tissue tension have identical effects on the mechanics of interphase and mitotic cells. (**B, D**) Box plots indicate the 25th and 75th percentiles, the red line indicates the median, and the whiskers extend to the most extreme data points that are not outliers. Individual data points are indicated by black dots and outliers by red crosses. (**A, C, E**) scale bars: 10 μm. **A.** Top: Representative profile of a dividing cell surrounded by interphase neighbouring cells within a monolayer at 0% strain, treated with DMSO or Y27632. Nucleic acids are visualised by H2B GFP (green), the cell membrane is labelled with CellMask 568 dye (white). Bottom: Diagram indicating apical angle of contact between the mitotic cell and its interphase neighbouring cells, Θ_*a,mi*_. **B.** Distribution of apical angles of contact between dividing cells and their interphase neighbours at 0% strain, measured in monolayers treated with DMSO or Y27632. WRST, p = 0.691. N=19 cells, DMSO; N=60 cells, Y27632. n=4 independent days, DMSO; n=8 independent experiments, Y27632. **C.** Top: Representative profile of a dividing cell surrounded by its interphase neighbouring cells within a monolayer at 0% strain, treated with DMSO. Yellow dashed lines indicate the profile of intercellular junctions around the mitotic cell or around interphase cells. Nucleic acids are visualised by H2B GFP (green), the cell membrane is labelled with CellMask 568 dye (white). Bottom: Diagram indicating measurement of the lateral radii of curvature of mitotic cells (red dashed lines). **D.** Distribution of lateral radii of curvature for dividing cells in monolayers at 0% strain, treated with DMSO or Y27632. WRST, p = 0.123. N=15 cells, DMSO;N=21 cells, Y27632. n=4 independent days, DMSO; n=8 independent days, Y27632. **E.** Relative surface tension in the apical, basal and junctional surfaces as a function of strain for control monolayers and monolayers treated with Y27632. Surface tensions are normalised to the basal tension for each strain magnitude. The error bar is the standard deviation of the monolayer mean surface tension (calculated as mean of 10 cells per monolayer). **F.** Absolute surface tension as a function of strain for apical, basal, and junctional surfaces for control and Y27632-treated monolayers. Basal tension is taken equal to the monolayer tension (Fig 3A), and apical and junctional tension are calculated using panel E. The error bar is the standard deviation of the monolayer mean surface tension (calculated as mean of 10 cells per monolayer).

As a complement to this analysis, we also measured the curvature of lateral junctions between mitotic cells and their interphase neighbours from XZ profiles, *R_l_*. This curvature reports on the difference in pressure between adjacent cells through Laplace’s law: 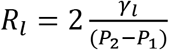, where *γ_l_* is the tension in the junction between two cells, and *P*_2_ and *P*_1_ are the pressures in each of the cells (**Fig 5C**). When intracellular pressure in adjacent cells is equal, their junction will appear straight (i.e. they have an infinitely large radius of curvature). Conversely, if the pressure is larger in one of the cells, the junction will curve away from the more pressurised cell. In suspended monolayers, junctions between interphase cells always appeared straight, whereas junctions between mitotic cells and interphase cells were always curved away from the mitotic cell. Changes in *R_l_* in response to a treatment would indicate that the pressure in mitotic and interphase cells is affected differentially. However, our experiments showed no change in curvature of lateral junctions, *R_l_*, when monolayers were treated with Y27632 (**Fig 5D**), again suggesting that interphase cells and mitotic cells are identically affected by Rho-kinase inhibition.

Overall, measurements of apical angle of contact, Θ_*a,mi*_, and curvature of lateral junctions, *R_l_*, indicate that, although mitotic cells are more contractile than interphase cells, the treatments used do not change the relative values of cortical tension and internal pressure between mitotic and interphase cells, nor dramatically change the morphology of the cells.

### Increased out-of-plane division correlates with low junctional tension

Our experiments could not identify morphological attributes associated with the alignment of cell division within the plane of the monolayer. However, tissue tension provided a more reliable predictor (**Fig 3C,D**). As astral microtubules and cortical contractility were both necessary to ensure accurate division in-plane, we hypothesized that spindles may sense tissue tension through its impact on tension at cellular interfaces relayed through astral microtubules. When they are integrated into epithelia, cells present clear differences in molecular composition, cytoskeletal organisation, adhesion, and signalling at their apical, lateral and basal surfaces as a result of the pathways that establish apico-basal polarity [43, 44]. As a result, the different subcellular surfaces likely differ in their mechanics. However, little is known about the relative magnitude of tension in each of these surfaces or how tissue-scale deformation affects each subcellular domain.

Assuming that cellular surfaces can be approximated to portions of sphere (see **Methods** and **Fig S7D**), the tension *γ* in a cellular surface is linked to its curvature r and the internal cellular pressure *P* through Laplace’s law. Therefore, measuring the apical and basal radii of curvature of interphase cells allows to determine relative changes in tensions in those surfaces as a function of strain (**Fig S7A**, see **Methods**). Indeed, Laplace’s law indicates that: 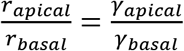. In addition, the ratio of junctional to apical tension can be inferred from geometrical and physical considerations as: 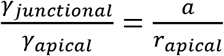 with *a* being the cell length (see **Methods**). Therefore, measuring *a, r_apical_*, and *r_basal_* as a function of applied strain allows to characterise the relative evolution of subcellular surface tensions in response to mechanical or chemical perturbations. We examined changes in radii of curvature in interphase cells because our modelling and experiments indicated that the mechanical changes in mitotic cells are proportional to those occurring in interphase cells and because this allows to gather sufficient data for statistical comparison between conditions.

At 0% strain, the radius of curvature of the basal side was approximately 5-fold larger than on the apical side (**Fig S7B, C**), indicating a larger tension on the basal surface that was consistent with the greater enrichment in myosin observed on the basal side [51, 45]. Both the mean apical and basal radii of curvature increased non-linearly with increasing strain, with the basal radius of curvature remaining systematically larger (**Fig S7B, C**). When we computed the relative tension at subcellular surfaces (normalizing basal tension to 1), we found that basal tension was higher than apical and junctional tensions – whose magnitude was very similar (**Fig 5E**). With increasing strain, both junctional and apical tension decreased relative to basal tension from ^~^0.3 at −30% strain to ^~^0.1 at 80% strain. As basal tension appeared several fold larger than apical tension, we approximated cellular basal tension to tissue tension (**Fig 3C**) to estimate absolute tensions in cellular surfaces. Using this approximation, basal tension grew linearly with strain from approximately 50 Pa at −30% strain to 950 Pa at 50% (**Fig 3C, 5F**). Apical and junctional tensions increased from 20-50 Pa at −30% strain to 100-200 Pa at 50% (**Fig 5F, Fig S7E,F)**. We found that apical and junctional tensions, respectively, were significantly lower at 0% strain in Y27632-treated monolayers than in controls, with magnitudes similar to those in control monolayers subjected to −30% strain (**Fig 5E, F**). The application of a 50% strain to Y27632-treated monolayers increased apical and basal tensions to levels comparable to controls at 0% strain (**Fig 5E, F**). These measurements together with our characterisation of the orientation of division for different applied tissue strains suggest that out-of-plane divisions are more frequent under conditions of low subcellular surface tension.

## Discussion

By mechanically manipulating suspended epithelial monolayers, we have demonstrated that increased frequency of out-of-plane divisions correlates with low tissue tension rather than with changes in cell shape or cell density. The impact of decreasing tissue tension was similar whether tension reduction was induced through the application of compressive strain or through chemical inhibition of myosin contractility. Furthermore, the accuracy of division orientations was restored using orthogonal means of returning tissue tension to similar levels as in control monolayers. Thus, the accuracy of in-plane divisions in monolayers subjected to compressive strain could be rescued by increasing cell contractility; while in monolayers in which contractility was inhibited, the frequency of out-of-plane divisions could be reduced by application of mechanical stretch. As our experiments and modelling indicate that tissue tension and cell surface tension evolve in similar ways, these data indicate that the molecular mechanism ensuring accurate in-plane division is sensitive to cellular surface tension but not the exact manner in which this tension is generated. Interestingly, our data are consistent with recent work showing that a sufficiently large cortical tension is necessary for spindles to orient along the cellular long axis in the drosophila notum midline [16]. While it is not clear precisely where subcellular surface tensions are read out by the spindle, our data suggest that junctional tensions are likely critical, since out-of-plane division was more frequent when astral microtubules were unable to contact the spindle orientation machinery present at intercellular junctions [22, 23, 13]. Indeed, in suspended monolayer epithelia, LGN localised to intercellular junctions and its localisation did not depend on strain or the activity of Rho-kinase. Thus, a sufficiently large junctional tension may be necessary to allow spindles to accurately orient cell division in the plane of the monolayer.

How cell surface tension influences the accuracy of spindle positioning in the plane is not known. In our experiments, we rarely observed significant repositioning of the spindle between metaphase and anaphase. Therefore, less accurate assembly and initial positioning out-of-plane is likely responsible for our observations. Centrosome and spindle positioning arises from a distribution of torques generated by interaction of forces between astral microtubules and the cortex [46, 20, 11]. Work in vitro has shown that both pushing and pulling forces contribute to aster centring [47, 48] and both could in principle be sensitive to cortical tension.

Pulling forces could be altered either by changes in localisation of cortical regulators or by changes in the pulling efficiency of dynein. Our data showed that localisation of cortical regulators was not changed in conditions that led to more out-of-plane divisions but that junctional tension decreased. Recent work has shown that spindles cannot orient along the long axis of cells if tissue tension is too low [16]. As our modelling shows that cell and tissue tension evolve in similar ways, this indicates that low cell surface tension likely impedes the ability of spindles to orient along the cell long axis. Therefore, we speculate that a less-tensed junctional cortex provides a less stable substrate upon which dynein motors can act to generate the pulling forces required to align spindles in the plane of the epithelium. In support of this, in *C. elegans* embryos whose F-actin cortex has been depolymerized, microtubules contacting the cell periphery extract membrane tethers rather than generate spindle centring forces [49]. Conversely, in adherent tissues, LGN recruitment to intercellular junctions was promoted by increased tension in E-cadherin complexes [14]. Thus, low tissue tension may decrease pulling forces arising from intercellular junctions through a combination of less LGN recruitment and less efficient pulling.

A role for pushing force appears less likely. Pushing forces are thought to arise from the microtubule plus end polymerising against the cortex. When the cortex is very stiff (or tensed), new GTP-bound tubulin heterodimers cannot be added to microtubules leading to hydrolysis of the GTP cap, catastrophes, and depolymerisation of the astral microtubules [48]. Conversely, when the cortex is less tensed, microtubule growth can continue by deforming the cortex, leading to an increase in pushing forces. In this scenario, because basal tension is several fold larger than apical tension, we would expect larger pushing forces to be generated by astral microtubules contacting the apical surface than the basal surface. In MDCK cells, astral microtubules emanate from the spindle poles with a large opening angle such that they likely contact each of the apical, basal, and junctional subcellular surfaces [31]. Therefore, we would expect that the larger pushing forces against the apical surface would displace spindles towards the basal side of cells. Since we did not observe this, our data suggest that the most likely cause of spindle misalignment in this system is the decrease in pulling forces induced by low junctional tension.

Out-of-plane cell divisions are observed in physiological contexts during stratification of multi-layered epithelia such as the skin and pathologically during hyperplasia and cancer. In all cases, multilayering is preceded by an increase in cell density which may decrease tissue tension. In monolayered epithelia, out-of-plane division orientation may act as a mechanism to maintain density homeostasis by retaining a single daughter cell. While it is not clear whether this change in the orientation of division arises from mechanotransduction monitoring tissue mechanics or is an emergent property of the forces driving spindle positioning, our data show that a decrease in tissue tension leads to a reduction in surface tension at the cellular scale that can cause division out of the plane because of a change in the forces exerted on astral microtubules. Thus, the initiation of multilayering may stem from a mechanical cue that is a natural consequence of the interaction between the spindle positioning machinery and the cortex of cells growing in epithelial monolayers under tension.

## Materials and methods

### Cell lines

MDCK cells were grown in DMEM (Thermo Fisher Scientific) supplemented with 10% fetal bovine serum (Sigma-Aldrich), HEPES buffer (Sigma-Aldrich), and 1 % penicillin/streptomycin in a humidified incubator at 37°C with 5% CO2. To visualise the DNA during division, cells were transduced with lentivirus encoding H2B GFP (kind gift from Dr Susana Godinho, Barts Cancer Institute, Queen Mary University of London, UK). To visualise the localisation of cortical pulling forces on astral microtubules, we generated a stable cell line expressing LGN-GFP. For this, LGN-GFP was excised from a plasmid (pTK14, plasmid #37360, Addgene) and inserted into pLPCX (Takara-Clontech). Retroviruses were then generated as previously described [51] and transduced into MDCK WT cells. All cell lines were selected with appropriate antibiotics and sorted by flow cytometry before use. Cells were routinely tested for the presence of mycoplasma using the mycoALERT kit (Lonza).

### Generating suspended MDCK monolayers

Suspended MDCK monolayers were made as described in [28]. Briefly, a drop of collagen was placed between two test rods and left to dry at 37C to form a solid scaffold. The dry collagen was then rehydrated and cells were seeded on top of it and cultured for 48-72 hours until cells covered the whole of the collagen and part of each test rod. Immediately before each experiment, the collagen scaffold was removed via collagenase enzymatic digestion, leaving the monolayer of cells suspended between the two test rods (**Fig 1A**).

### Imaging suspended MDCK monolayers

Tissues were imaged at 37C in a humidified atmosphere with 5% CO2. The imaging medium consisted of DMEM without phenol red supplemented with 10 % FBS. To visualize the shape of the cells during division, cell membranes were labelled with CellMask membrane stain for 10 min prior to collagen digestion following the manufacturer protocol (Thermo Fisher Scientific). To visualise the boundaries of the suspended monolayer, AlexaFluor-647-conjugated dextran, 10,000MW (Thermo Fisher Scientific) was added at 20 μg ml^-1^ to the imaging medium.

XYZ stacks of the tissue before and after mechanical manipulation were obtained using a 40x objective (UPlanSApo, 1.25 Sil), on an Olympus IX83 inverted microscope equipped with a scanning laser confocal head (FV-1200, Olympus, Berlin, Germany). A single stack was obtained before, and the tissue was imaged for approximately 1h after the mechanical manipulation, acquiring stacks at intervals of 1-2 minutes with Z-slices spaced by 1 μm.

### Mechanical manipulation of suspended MDCK monolayers

Mechanical manipulation of the MDCK monolayers along the X-axis was performed as previously described [29]. A custom-made steel wire probe was connected to a 2D manual micromanipulator which was mounted onto a motorized platform (M-126.DG1 controlled through a C-863 controller, Physik Instrumente, Germany). The manual micromanipulator was used to position the probe so that it was wedged into a ‘V’-shaped section of the stretching device arm, allowing both forward and backward movement to compress and stretch the monolayer (**Fig 1A**). The tissues were deformed by controlling the motion of the motorized platform with a custom-written LabView program (National Instruments, USA). To ensure that the same part of the tissue was imaged before and after mechanical manipulation, the microscope stage (PS3J100, Prior Scientific Instruments, USA) was moved such that it matched the movement of the motorised platform using our LabView program to synchronise motion.

To apply cyclic stretch, our LabView programme generated sinusoidal displacement of the required amplitude and period.

### Stress-strain relationships in suspended MDCK monolayers

To measure the evolution of strain applied at the tissue level, the entire width of the tissue was imaged with a 4x objective and bright field illumination at 1 second intervals on an inverted microscope with environmental control. To extract the strain exerted on the tissue from these videos, a script (Mathematica, Wolfram, USA) was written which used a Hough transform to detect the edges of the stretching device to which the tissue was attached. This data was used to compute the change in length of the tissue and the tissue strain.

Tissue-scale stress was varied by moving one of the test rods with the motorised micromanipulator (see **Fig 3A**). Then, tissue stress was measured as described in [32]. Briefly, one extremity of the tissue was connected to a Nitinol wire which served as a force cantilever thanks its shape-memory properties. The force *F* exerted by the tissue was then deduced from the deflection *d* of the wire with respect to its reference position x_0_: *d* = *x* − *x*_0_ and *F* = *k.d*, with *k* the stiffness of the wire and *x* the position of the wire. These positions were extracted from images of the monolayer acquired using a 2X objective (Olympus) mounted on an inverted microscope (Olympus IX-71) equipped with a CCD camera (GS3-U3-60QS6M-C, Pointgrey). To define the reference position of the wire *x_0_*, tissues were detached from the device by cutting them with a tungsten needle at the end of the experiment. Stress was then defined as 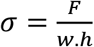 with *w* the average width of the tissue and *h* the tissue thickness. Ramps of strain were applied at a strain rate of 0.1%.s^-1^, a rate at which tissues exhibited a purely elastic behaviour with no viscous contribution [32]. Tissue-scale strain was extracted as described above.

### Quantification of cell strain during monolayer deformation

To measure strain at the cell level, MDCK cells labelled with CellMask membrane stain were imaged using a 60x oil immersion objective. A region of the monolayer, which contained both mitotic and interphase cells, was chosen close to the rigid arm of the stretching device. This region was maintained in the field of view of the camera during cyclic stretch and time-lapse imaging via manual movements in XY using the motorised microscope stage and refocusing. A custom-written script (Mathematica) used image cross-correlation to align the frames of the resulting videos and the Tissue Analyzer plugin of Image J was used to segment the cell shapes. Bounding boxes were fitted to each cell at each time point to calculate the temporal evolution of cell strain.

### Quantification of cell shape and orientation in the XY-plane

By convention, the X-axis was taken as the axis of deformation and the Y-axis was perpendicular to that. The shape of mitotic and interphase cells was characterised from confocal microscopy image stacks of monolayers stained with CellMask. The cell shape was manually marked using Fiji (http://fiji.sc/Fiji). The ellipse that best fitted the cell outline was calculated in Fiji; the length of the bounding box of the ellipse along the X-axis was taken as a measure of cell size along the X-axis, the width of the bounding box of the ellipse was taken as the cell size along the Y-axis, and the ellipse orientation was used as a measure of cell orientation. Measurements of cell height (cell size along the Z-axis) were obtained manually in Fiji from sectioning of the image stack along XZ planes. Cell shape and orientation were determined in the XYZ stack before and immediately after mechanical manipulation and additionally in the XYZ stack at the beginning of metaphase.

### Quantification of spindle orientation out-of-plane

Spindle orientation with respect to the Z-axis was determined at the beginning of metaphase, when the metaphase plate was formed, (Θ_*m*_) and at the end of division, when closing of the cytokinetic furrow was complete, (Θ_*d*_). The angles were measured from the cross-sectional sections through the image stack along XZ planes. The metaphase spindle orientation (Θ_*m*_) was measured as the angle between the line going through the metaphase plate and the line perpendicular to the monolayer plane (**Fig 1D**). Spindle orientation at the end of division (Θ_*d*_) was measured as the angle between the line going through the new junction between daughter cells and the line perpendicular to the monolayer plane (**Fig 1D**).

### Quantification of metaphase plate length and spindle length

Spindle length and metaphase plate length were measured at the beginning of metaphase from the cross-sections through the image stack along XZ planes. To visualise the spindle, MDCK H2B GFP cells were incubated for 30 min before the start of imaging with the SiR-tubulin dye (Spirochrome, Switzerland). Spindle length was measured manually in Fiji as the distance between the two spindle pole bodies. Metaphase plate length was determined from the H2B GFP signal.

### Quantification of cellular radii of curvature

Apical, basal and lateral mitotic cell outlines were manually marked from the CellMask signal at the beginning of metaphase in cross-sections through the image stack along XZ planes using Fiji. A circle was fitted through each of the marked outlines, and the respective radii were tabulated. Similarly, apical and basal outlines of interphase cells were manually marked from the CellMask signal and radii of curvature were determined in the same way as for the mitotic cells.

### Quantification of apical contact angle

The apical contact angle, *θ_a,mi_*, between mitotic cells and their neighbours was measured at the beginning of metaphase from the CellMask signal in cross-sections through the image stack along XZ planes using Fiji.

### Drug treatments

To block Rho-kinase activity, monolayers were treated with Y-27632 (Tocris, UK) at a concentration of 50 μM, 10 minutes prior to imaging. To inhibit polymerisation of astral microtubules without significantly affecting the spindle, monolayers were treated with low doses of nocodazole (20 nM, Merckmillipore, UK) for 10mins before experiments. To increase myosin contractility, myosin phosphatases were inhibited by addition of 35 nM of calyculin A (Sigma-Aldrich, USA), 10 min before experiments. All drugs were dissolved in DMSO.

### Tubulin immunostaining

For immunostaining of microtubules, cells were fixed in ice-cold methanol at −20 before washing three times in PBS containing 10% horse serum (HS) for 5min each and being incubated with a mouse monoclonal primary antibody against α-tubulin (DMA1, 1:1000 dilution, Abcam, UK) for 1 hour at RT. This was followed by three washes in PBS+10%HS for each lasting 5min, incubation in goat anti-mouse secondary antibody conjugated with Alexa 488 (Thermofisher, 1:200 dilution) for 1hour at RT, three washes in PBS+10%HS each lasting 5min and staining of nucleic acids with Hoechst 33342 (5μg/mL-Merck Bio-sciences) for 5min. Following staining, cells were mounted in FluorSave reagent (Merckmillipore, UK) and imaged on Olympus FV-1200 using a 100x objective (NA 1.40, Olympus, Germany).

### Estimation of relative tension at subcellular surfaces

We use geometrical considerations to estimate the evolution of apical, lateral, and basal tensions from experimental measurements of the apical and basal radii of curvature at different tissue strains using Laplace’s law.

Because Laplace’s law assumes that surfaces can be approximated to portions of spheres, we first verified that monolayer profiles were similar when viewed along the axis of deformation (xz profiles) and perpendicular to it (yz profiles). To verify this, we measured the radii of curvature of interphase cells along profiles acquired in the xz and yz directions and found that these were not significantly different on either the apical or the basal side (**Fig S4F**). Based on these measurements, we concluded that cell apices and bases could be approximated to portions of a sphere.

According to Laplace’s law, the radius of curvature is dictated by the interplay between the internal cellular pressure *P* and the surface tension : 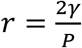. As apical and basal surfaces are exposed to the same internal pressure, the ratio of surface tensions can simply be inferred from the ratio of radii of curvature: 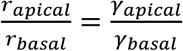.

At the apical surface, we can use the Young-Dupré relationship to estimate *Y_junctional_* from *γ_apical_*:

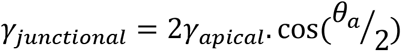

with *θ_a_* the apical angle of contact. If we approximate the apical side of the monolayer to a series of portions of circle with the same radius of curvature *r_apical_* and assume that all cells have a width *a* (**Fig 5E**), *θ_a_* can be estimated as:

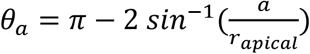

By combining the two equations, we obtain:

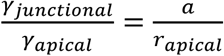

Similar relationships can also be written for the relationship between junctional and basal tensions. In this study, we normalise tensions to *γ_basal_* because our previous work has shown that tension on the basal surfaces is highest (45).

Therefore, by measuring *a, r_apical_*, and *r_basal_* as a function of applied strain, we can determine how surface tensions vary with respect to one another with strain.

### Computational modelling

To mimic experimental testing of suspended monolayers, we use a finite element model implemented in Julia (**Fig S4**). In this model, the tissue is discretised as a triangular mesh where each cell is represented by a hexagon consisting of six triangles. Each triangle side is modelled as a spring with spring constant *k*, length *l*, and rest length *l*_0_. The mechanical connection between springs is modelled with pins that allow free rotation around the extremities of each bar.

We assumed that the stress-free shape of the monolayer was a rectangle of 20 x 20 hexagonal cells. To simulate myosin contractility within each cell, we imposed an initial prestress *γ* in each spring by setting the rest length *l_0_* such that *γ* = *k*(*l_init_* − *l*_0_), where *l_init_* is the length in the reference configuration (0% strain). To mimic the presence of mitotic cells within the monolayer, we modified the stiffness *k* and the contractility *γ* of one hexagon within the centre of the monolayer such they were 2.5-fold and 2-fold larger, respectively, to match the stiffer spring constant and greater contractility observed in mitotic cells (blue elements, **Fig S4**, **Fig 4B**).

Displacement boundary conditions were imposed to nodes on each of the vertical edges of monolayer to allow simulation of uniaxial deformation and the horizontal edges were left free (**Fig S4**). To simulate compressive/tensile loading, displacements *d* were imposed along the horizontal axis (x-axis). To identify the equilibrium configuration, we used the finite element method [52].

From the displacements of each node, we can compute the stress tensor acting on any subarea *A* as

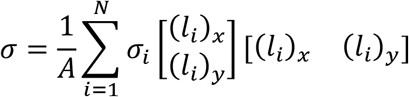

where *N* is the number of elements within the sub-area, *σ_i_* is the stress along element *i*, and *l_x_* and *l_y_* are the length of element *i* projected along the x- and y-axis, respectively.

The model is first used to assess the stress at different cell locations assuming no mitotic cells are present. By comparing the principal stresses along the x- and y-directions close to the centre of the monolayer to those occurring further away from the centre (thus comparing the stress tensors in columns 2 vs 4 and 3 vs 5 in **Fig S5**), we conclude that the stress tensors within the monolayer are approximately homogeneous, consistent with the experimental observation of homogeneous strain in stretched monolayers [51]. Therefore, the stress on the test rods measured in experiments is a good approximation of the stress experienced by cells in the centre of the monolayer. A similar conclusion was reached when a dividing cell is present in the centre of the monolayer (**Fig S6**).

**Table 1:** Model parameters for simulation of different experimental conditions displayed in **Figs S5, S6**.

| Simulation | Spring constant k | Strain | Contractility $\gamma$ |
| --- | --- | --- | --- |
| Control – Untreated monolayer | 1 | 0 | 0.35 |
| Compressed untreated monolayer | 1 | -30% | 0.35 |
| Compressed monolayer with increased contractility | 1.5 | -30% | 0.5 |
| Monolayer with reduced contractility | 0.7 | 0 | 0.1 |
| Stretched monolayer with reduced contractility | 0.7 | 50% | 0.1 |

### Statistical and data analysis

All other data and statistical analysis was performed in MATLAB R2018a (Mathworks, Natick, MA, USA).

Boxplots show the median value (red line), the first and third quartile (bounding box) and the range (whiskers) of the distribution. The red crosses mark the outliers. Raw data points are plotted on top of all boxplots. All tests of statistical significance are Wilcoxon rank-sum tests unless otherwise stated.

Fischer exact tests were used to assess the change in the proportion of cells dividing in plane in response to a treatment. For this, we categorised dividing cells as having low Z-angles (<30°, in-plane) or high Z-angles (>30°, out-of-plane). We then computed the Fischer exact test statistics using SPSS (IBM, Armonk, NY, USA).

Image processing was performed with Fiji.

## Data availability

All reagents are available from the corresponding author upon request.

Code and primary data are available from the UCL data repository (https://rdr.ucl.ac.uk/) with a unique doi (10.5522/04/16930864).

## Acknowledgements

We thank Maxine Lam, Susana Godinho (Queen Mary University of London, London, UK), and past and present members of the Charras and Baum labs for comments and discussions. AL was supported by an EMBO Long-term fellowship (ALTF 29-2016). MK was supported by a SNSF early post-doc fellowship P2LAP3_164919. AL, MK, JD, TW, and JF were supported by a European Research Council consolidator grant (CoG-647186) to GC.

## Author information

AL and GC conceived the project and wrote the manuscript. AL did most of the monolayer experiments and performed the analysis. JF performed monolayer stress measurements, contributed to discussions and analysis. MK performed tubulin immunostainings, contributed to experiments and discussions. TW performed measurements of relative stiffness of mitotic and interphase cells. JD performed nocodazole monolayer experiments. ABN performed measurements of apical contact angles. AB and AK devised and implemented the theoretical model. BB advised on experiments and discussions. GC oversaw the entire project.

## Supplementary Figures

**Figure S1.**
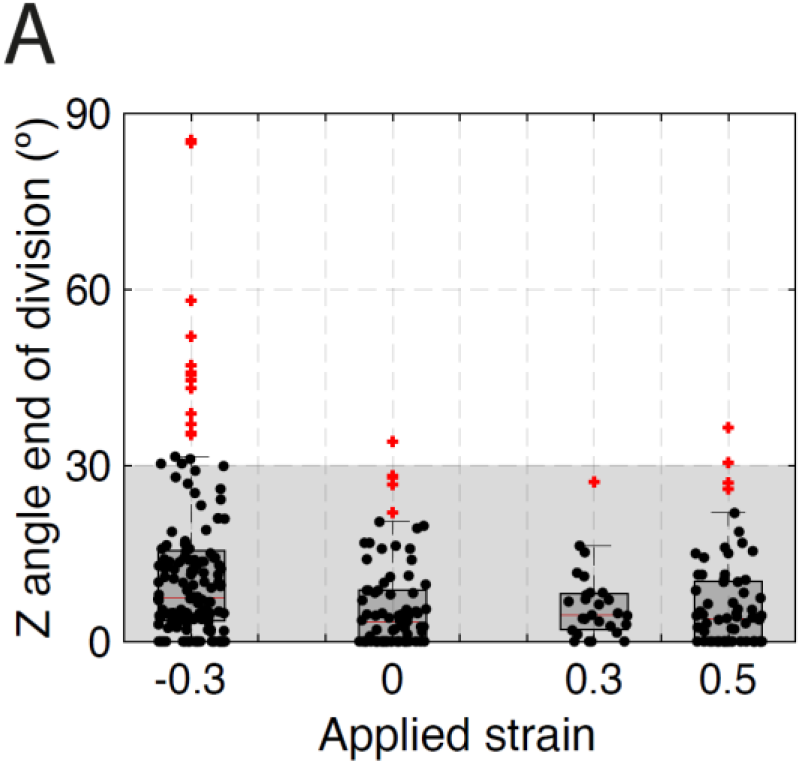
related to Figure 1. Z angle at the end of division for different applied deformations. **A.** Distribution of the spindle Z angle at the end of division for different applied strains in control monolayers. Box plots indicate the 25th and 75th percentiles, the red line indicates the median, and the whiskers extend to the most extreme data points that are not outliers. Individual data points are indicated by black dots and outliers by red crosses. Gray box highlights Z angles < 30º. The number of cells examined for each condition was N=156 for −30% strain, N=80 for 0% strain, N=27 for 30% strain, and N=69 for 50% strain. Experiments were performed on n=14, n=8, n=4, and n=8 independent days, respectively.

**Figure S2.**
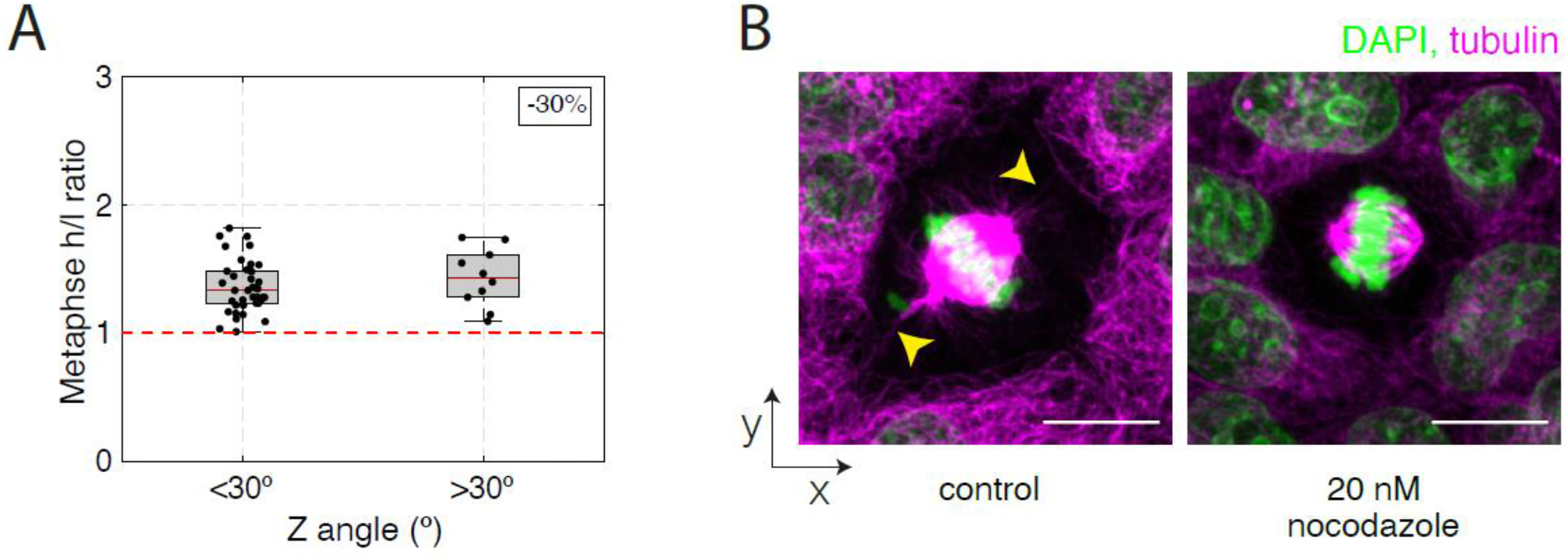
**related to Figure 2. A.** Ratio of cell height/length at metaphase as a function of spindle Z angle at metaphase for dividing cells in compressed monolayers (−30 %). Metaphase spindle Z-angles were categorised as either in-plane (Θ<30°), or out-of-plane (Θ >30°). Data as in Fig 2D. Box plots indicate the 25th and 75th percentiles, the red line indicates the median, and the whiskers extend to the most extreme data points that are not outliers. Individual data points are indicated by black dots and outliers by red crosses. **B.** Representative immunofluorescence images of WT MDCK cells stained for α-tubulin. Cells were treated with DMSO (control) or with 20nM nocodazole to disrupt astral microtubules. Yellow arrows indicate astral microtubules. DNA is stained with DAPI and shown in green, tubulin staining is shown in magenta. Scale bar: 10μm.

**Figure S3.**
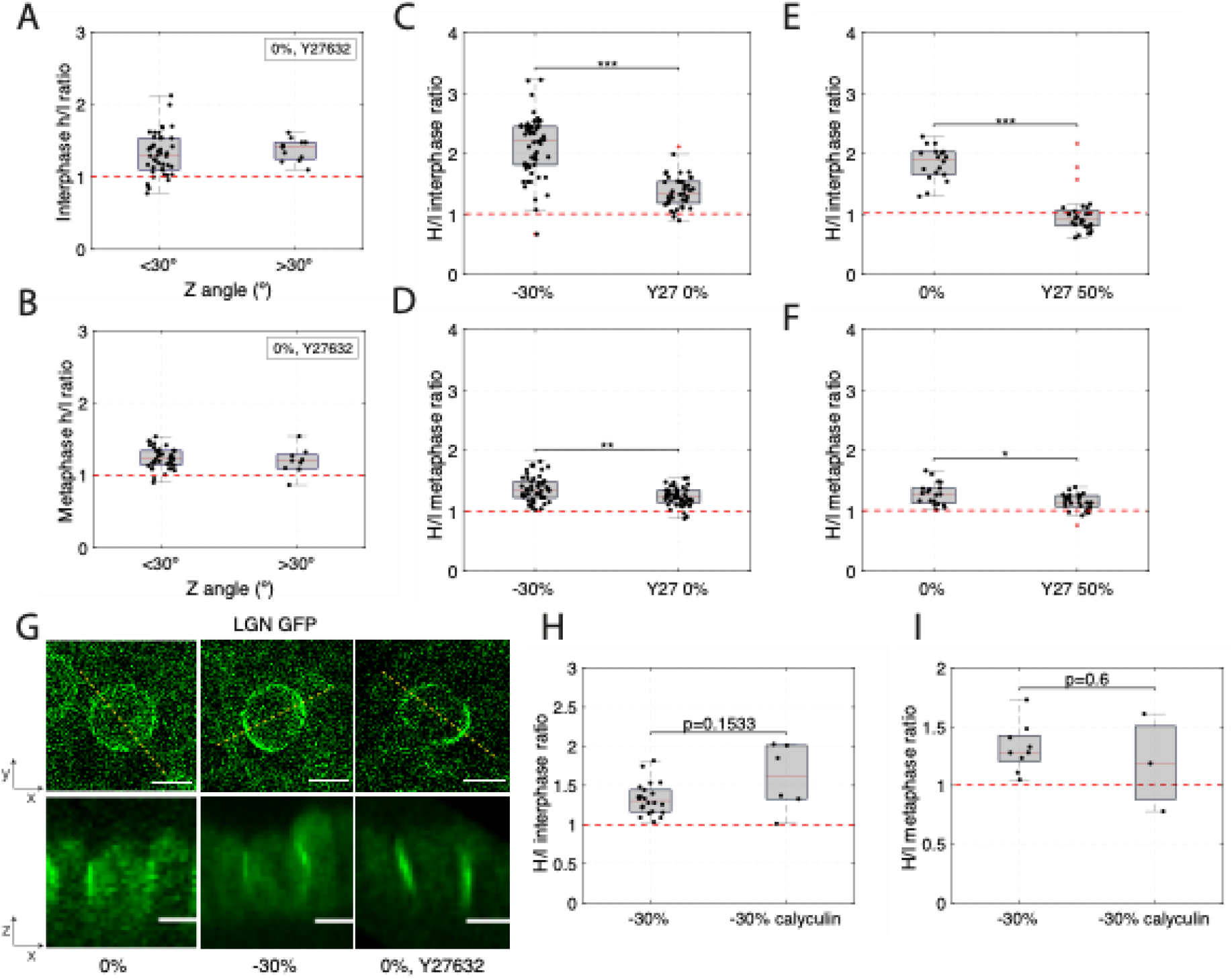
related to Figure 2. Cell shape and the localisation of spindle positioning proteins do not correlate with poor in-plane orientation. **(A-F, H-I)** Box plots indicate the 25th and 75th percentiles, the red line indicates the median, and the whiskers extend to the most extreme data points that are not outliers. Individual data points are indicated by black dots and outliers by red crosses. **A.** Height/length ratio of interphase cells in monolayers subjected to 0% strain and treated with Y27632 as a function of metaphase Z angle from dividing cells. Metaphase spindle Z-angles were categorised as either in-plane (Θ<30°), or out-of-plane (Θ >30°). Data from Fig 3D, E. **B.** Height/length ratio of metaphase cells in monolayers subjected to 0% strain and treated with Y27632 as a function of metaphase Z angle from dividing cells. Metaphase spindle Z-angles were categorised as either in-plane (Θ<30°), or out-of-plane (Θ >30°). Data from Fig 3D, E. **C.** Height/length ratio of interphase cells in monolayers subjected to 30%compressive strain or to 0% strain and treated with Y27632. WRST, p = 1×10^-11^. Data from Fig 1E and Fig 3E. **D.** Height/length ratio of metaphase cells in monolayers subjected to 30% compressive strain or to 0% strain and treated with Y27632. WRST, p = 0.002. Data from Fig 1E and Fig 3E. **E.** Height/length ratio of interphase cells in monolayers at 0% strain or treated with Y27632 and subjected to 50% strain. WRST, p = 3×10^-7^. Data from Fig 1E and Fig 3E. **F.** Height/length ratio of metaphase cells in monolayers at 0% strain or treated with Y27632 and subjected to 50% strain. WRST: p = 0.017. Data from Fig 1E and Fig 3E. **G.** Representative localisation of LGN in dividing cells in a monolayer subjected to 0 % strain, −30% compressive strain, and 0% strain with Y27632 treatment viewed in the XY (top) and XZ (bottom) planes. Dashed yellow lines indicate the locations at which XZ profiles were taken. Scale bars: 10 μm. **H.** Height/length ratio of interphase cells in monolayers subjected to 30% compressive strain, with and without calyculin treatment. WRST, p = 0.15. Data from Fig 3G. **I.** Height/length ratio of metaphase cells in monolayers subjected to 30% compressive strain, with and without calyculin treatment. WRST, p = 0.6. Data from Fig 3G.

**Fig S4:**
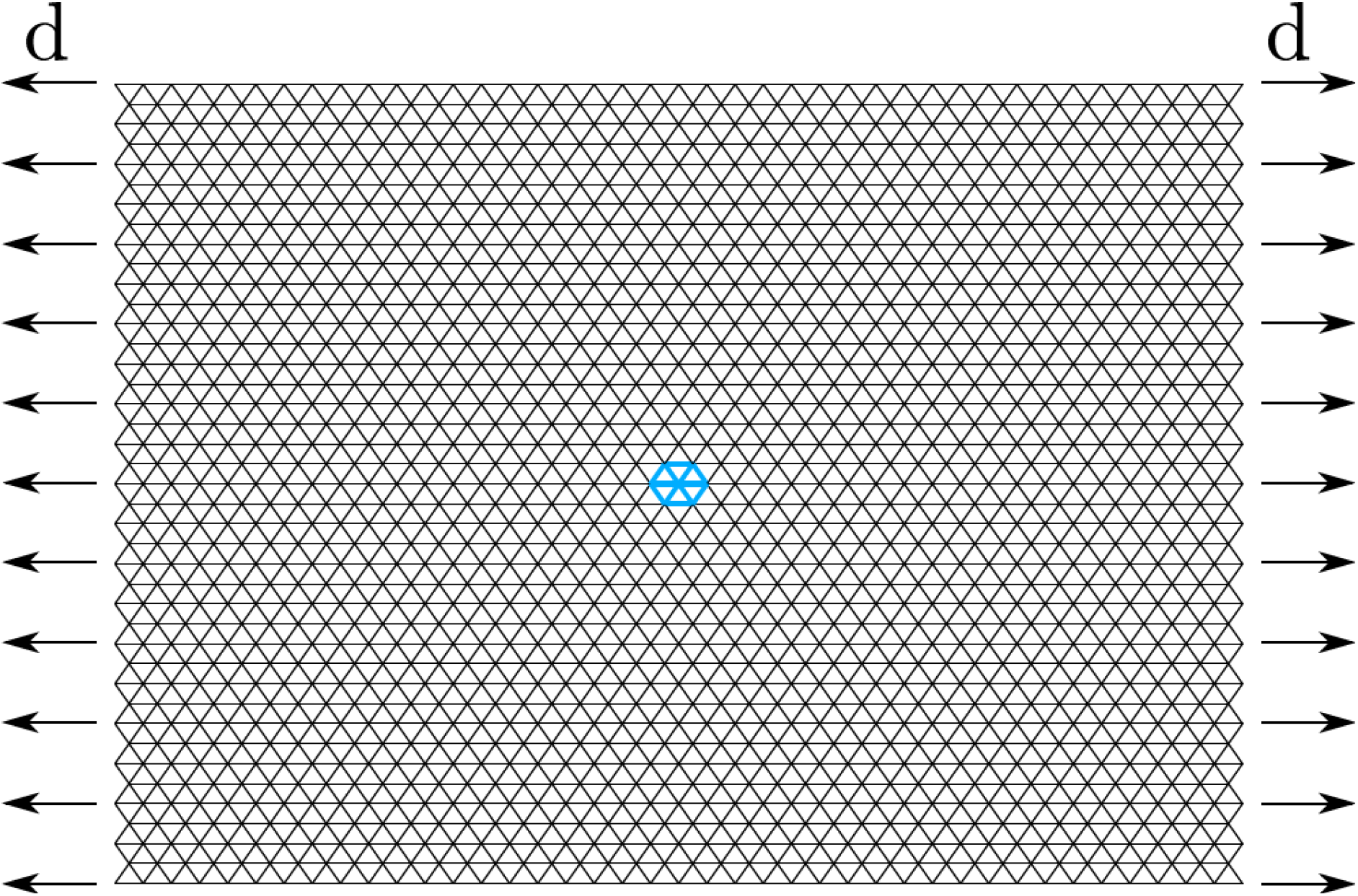
Sketch of the 2D planar monolayer model. The epithelial monolayer is discretised into triangles and subjected to displacement boundary conditions at its vertical edges while the horizontal edges are left free. Each triangle edge consists of a linear elastic element. The blue elements at the centre of the structure exemplify a mitotic cell (i.e. the stiffness and contractility of these structural elements are consistently increased to simulate a dividing cell).

**Fig S5:**
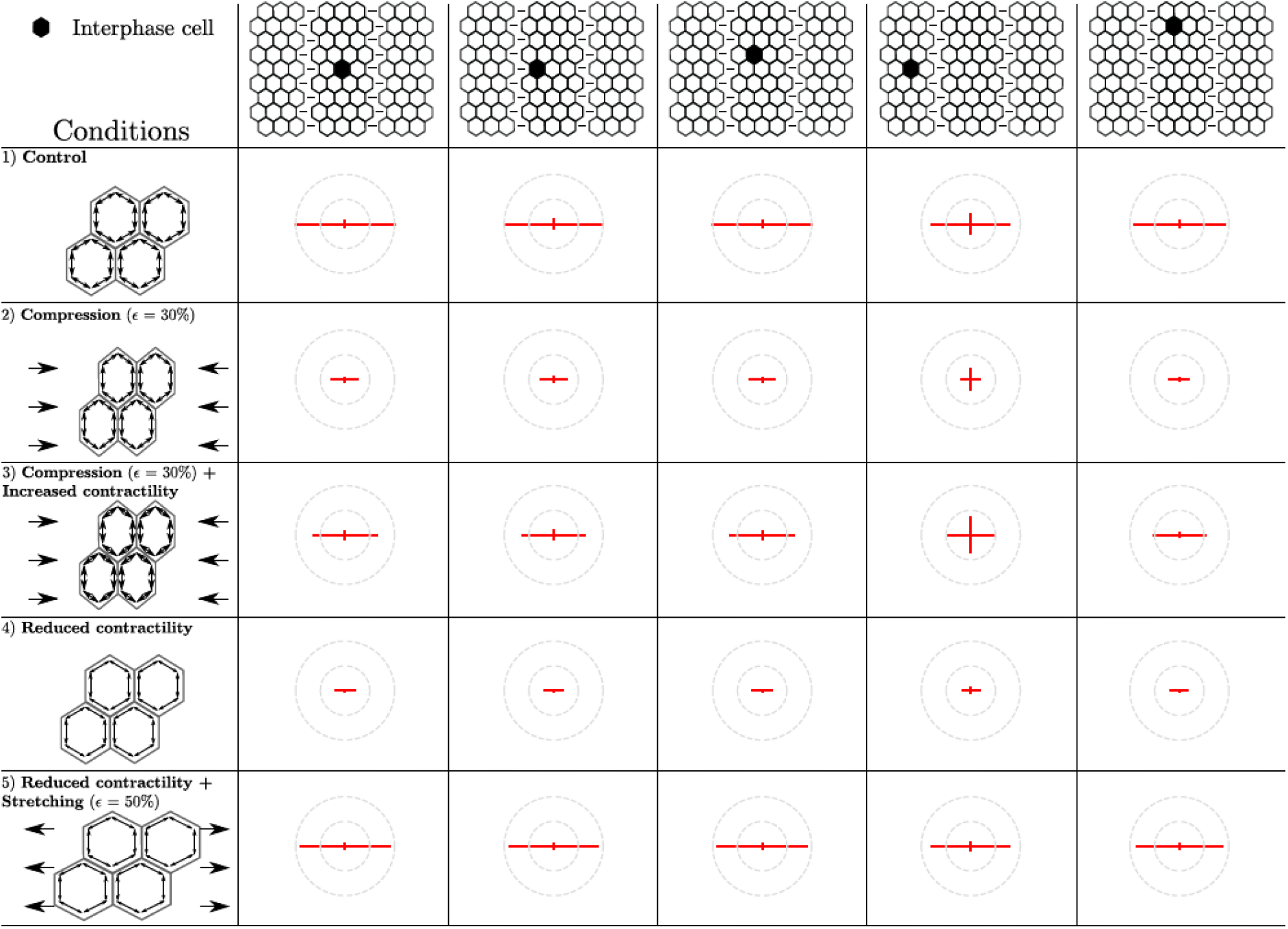
Principal stresses in cells within monolayers consisting entirely of interphase cells. Principal stresses are represented by orthogonal line segments with a length equal to the stress amplitude and a colour encoding their sign (red for tensile stress and blue for compressive stress). The stress tensor is plotted for different cell locations (each corresponding to a column): a cell at the centre of the monolayer, a cell directly adjacent to the central cell, a cell directly above the central cell, a cell far from the central along the x-axis, and a cell far above the central cell on the y-axis. Each row depicts a different experimental condition: untreated monolayer, compressed monolayer, compressed monolayer with increased contractility (via calyculin treatment), unloaded monolayer with reduced contractility (via Y27632 treatment), stretched monolayer with reduced contractility. Stress tensors are normalised to the stress tensor in control conditions (0% strain).

**Fig S6:**
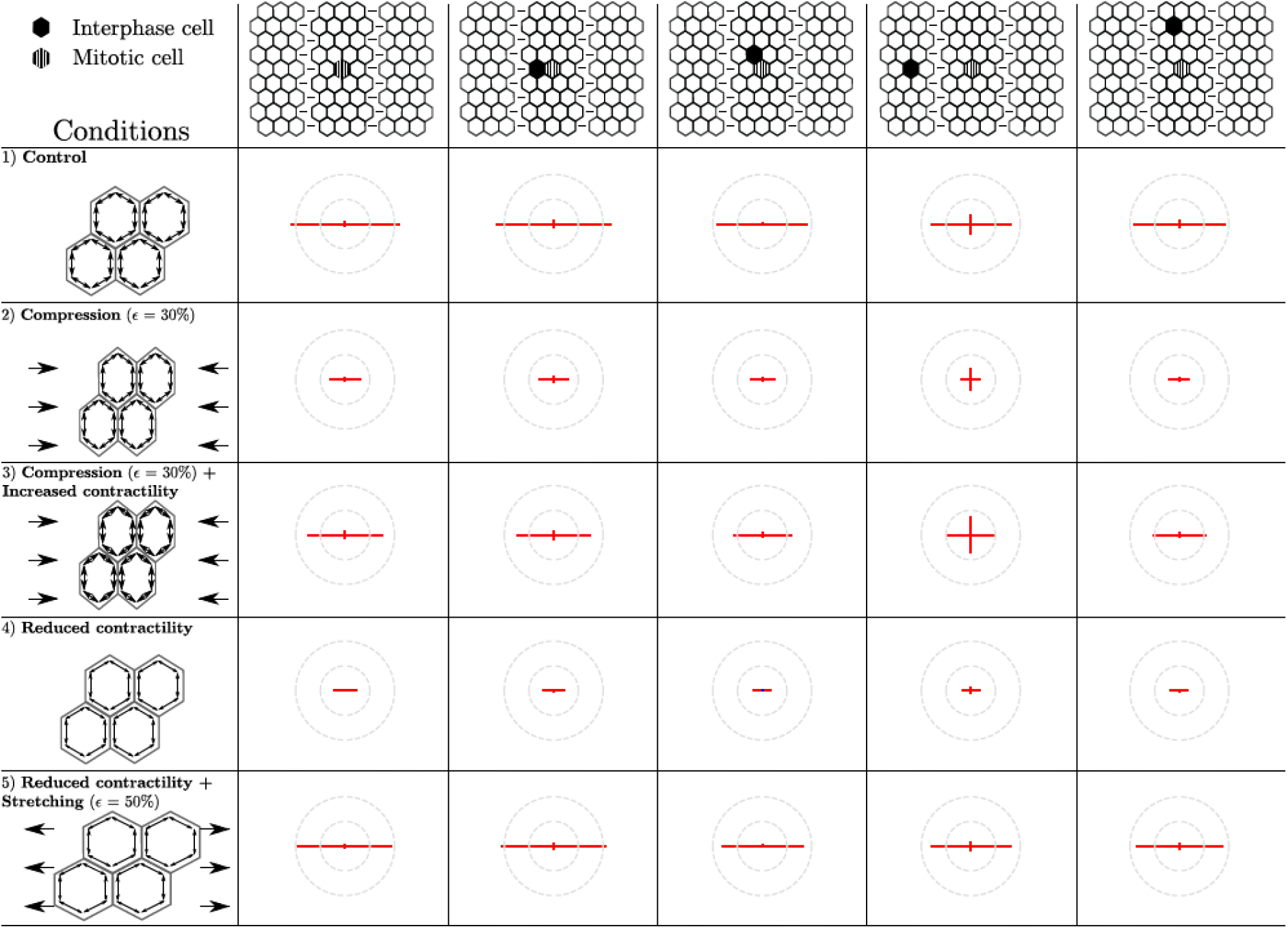
Principal stresses in cells within monolayers consisting of interphase cells with a mitotic cell in their centre. The mitotic cell is represented by the hashed hexagon. Principal stresses are represented by orthogonal line segments with a length equal to stress amplitude and a colour encoding their sign (red for tensile stress and blue for compressive stress). The stress tensor is plotted for different cell locations (each corresponding to a column): the mitotic cell in the centre of the monolayer, an interphase cell directly adjacent to the mitotic cell, an interphase cell directly above the mitotic cell, an interphase cell far from the mitotic cell along the x-axis, and an interphase cell far above the mitotic cell on the y-axis. Each row depicts a different experimental condition: untreated monolayer, compressed monolayer, compressed monolayer with increased contractility (via calyculin treatment), unloaded monolayer with reduced contractility (via Y27632 treatment), stretched monolayer with reduced contractility. Stress tensors are normalised to the stress tensor in control conditions in the centre of the monolayer without inclusion (**Fig S6**, first row, first column).

**Figure S7.**
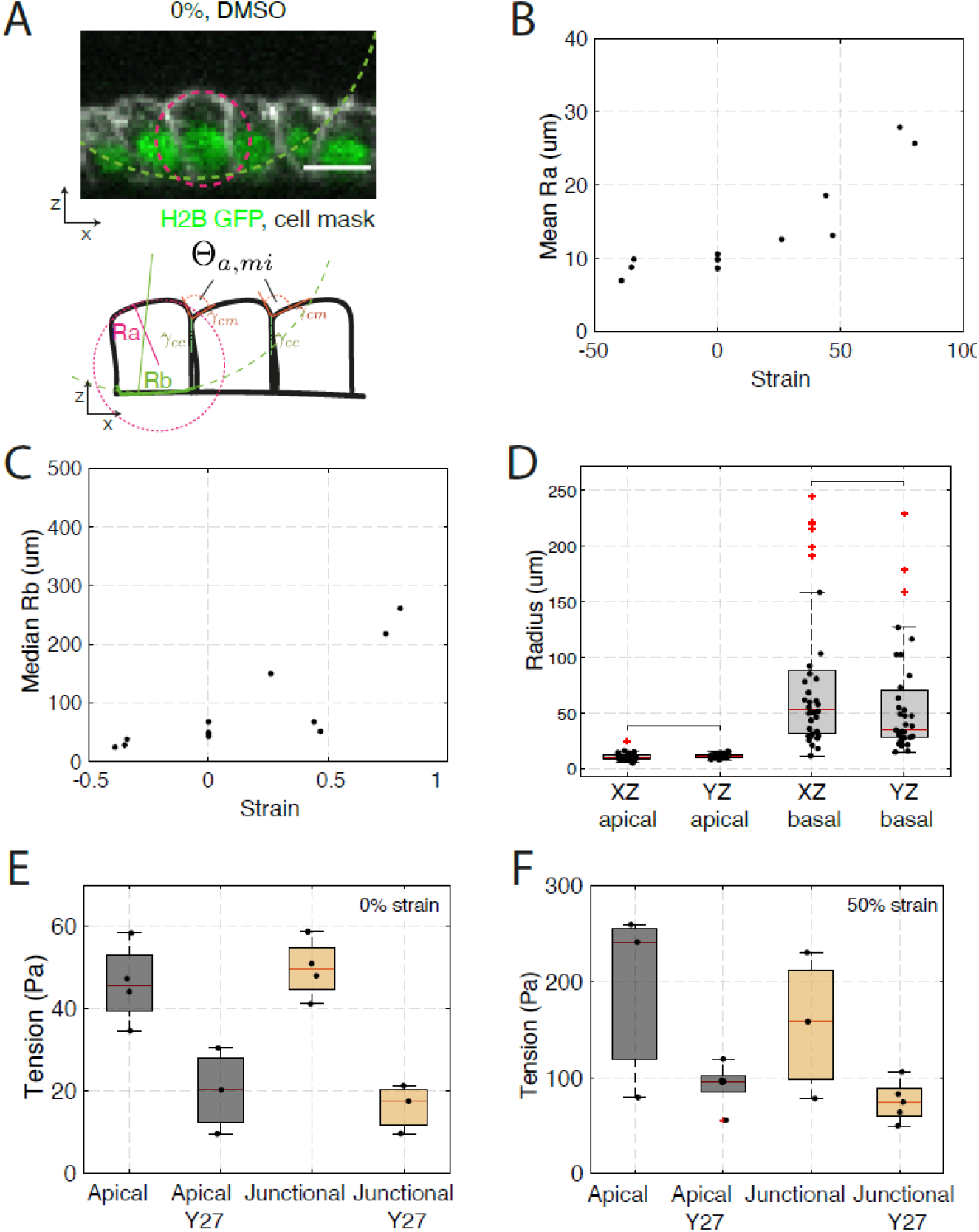
The apical and basal radii of curvature increase non-linearly with strain. **A.** Top: Representative profile of interphase cells within a monolayer at 0% strain, treated with DMSO. The basal and apical radii of curvature are indicated by a green dashed line and a magenta dashed line, respectively. Nucleic acids are visualised by H2B GFP (green), the cell membrane is labelled with CellMask 568 dye (white). Bottom: Diagram of the profile of interphase cells indicating the apical contact angles, *θ_a_*, the tension at the intercellular junctions *γ_junctional_* and the tension at apical junctions *γ_apical_* (**see methods**). **B.** Mean apical interphase radius as a function of strain. Each data point represents the average apical radius from a minimum of 20 interphase cells in a monolayer. Values for three monolayers are given for each strain range. **C.** Median basal interphase radius as a function of strain. Each data point represents the median apical radius from a minimum of 20 interphase cells in a monolayer. Values for three monolayers are given for each strain range. **D.** Apical and basal radii measured in the XZ and XY direction. Each data point represents one cell. A minimum of 10 cells per monolayer from 3 different monolayers at 0% strain were measured. Box plots indicate the 25th and 75th percentiles, the red line indicates the median, and the whiskers extend to the most extreme data points that are not outliers. Individual data points are indicated by black dots and outliers by red crosses. No significant differences were detected between radii measured in the XZ and YZ directions. **E.** Absolute surface tension of apical and junctional surfaces for control and Y27632-treated monolayers at 0% strain (as in Fig 5F). **F.** Absolute surface tension for apical and junctional surfaces for control and Y27632-treated monolayers at 50% strain (as in Fig 5F).

